# An atlas and database of neuropeptide gene expression in the adult zebrafish forebrain

**DOI:** 10.1101/2023.03.29.534505

**Authors:** Towako Hiraki-Kajiyama, Nobuhiko Miyasaka, Reiko Ando, Noriko Wakisaka, Hiroya Itoga, Shuichi Onami, Yoshihiro Yoshihara

## Abstract

Zebrafish is a useful model organism in neuroscience; however, its gene expression atlas in the adult brain is not well developed. In the present study, we examined the expression of 38 neuropeptides, and glutamatergic neuron marker gene mix (*slc17a6a, slc17a6b, slc17a7a,* and *slc17a7b*) in the adult zebrafish brain using *in situ* hybridization. The results are summarized as an expression atlas in 19 coronal planes of the forebrain. Furthermore, the scan data of all sections were made publicly available as a database. Based on these data, we performed detailed neuroanatomical analyses of the hypothalamus. By scrutinizing and comparing the expression patterns of neuropeptides, we found that several regions described as one nucleus in the reference zebrafish brain atlas contain two or more subregions with significantly different neuropeptide/neurotransmitter expression profiles, and we proposed them as novel subnuclei. Subsequently, the expression data obtained in this study were compared with those in mice, and a cluster analysis was performed to examine the similarities. As a result, several nuclei in zebrafish and mice were clustered in close vicinity: zebrafish ventral part of the anterior part of the parvocellular preoptic nucleus (PPav)/magnocellular preoptic nucleus (PM) and mouse paraventricular hypothalamic nucleus (Pa), zebrafish posterior part of the parvocellular preoptic nucleus (PPp) and mouse medial preoptic area (MPA), zebrafish dorsal part of the ventral zone of periventricular hypothalamus (Hvd)/anterior tuberal nucleus (ATN) and mouse ventromedial hypothalamic nucleus (VMN). The present expression atlas, database, and anatomical findings will contribute to future neuroscientific research using zebrafish.

**Key points:**

- The expression of 38 neuropeptides and GABAergic/glutamatergic neuronal marker genes in adult zebrafish forebrain was examined and compiled as an atlas.
- All scanned brain section data were published as a database.
- Based on the expression data obtained, multiple subnuclei in the zebrafish hypothalamus were proposed, and comparisons with the mouse hypothalamus were conducted.

## 1. Introduction

Zebrafish (*Danio rerio*) is widely used as a vertebrate animal model in various biology research fields. It has several advantageous properties, such as external fertilization, large clutch size, transparency of eggs and larvae, rapid development, and feasibility for both forward and reverse genetics (Kalueff *et al*., 2014; Stewart *et al*, 2014; Gerlai, 2020). Recently, whole-brain calcium imaging and high-throughput behavioral assays have been successively developed for larval zebrafish, leading to breakthroughs in neuroscience research (Orger and Polavieja, 2017; Vanwalleghem *et al*., 2018; Loring *et al*., 2020; Lee *et al*., 2022). In addition, multiple brain databases have been developed for larval zebrafish, such as the gene expression atlas of 2-, 3-, and 4-days post-fertilization (dpf) larval brains (ViBE-Z, Ronneberger *et al*., 2012), the brain atlas of single neuron tracings and images of transgenes and *in situ* hybridizations in 6-dpf larvae (MapZebrain, Kunst *et al*., 2019), and transgene expression- and immunohistochemistry-based atlas in 6-dpf larvae (Z-Brain Atlas, Randlett *et al*., 2015; Zebrafish Brain Browser 2 (ZBB2), Tabor *et al*., 2019). However, the number of available brain database resources on adult zebrafish is limited so far, which include the reference brain atlas (Wullimann *et al*., 1996), a database of transgene expression patterns in enhancer-trap lines (ZeBrain), magnetic resonance histology-based 3D atlas (Ullmann *et al*., 2010), and 3D atlas of immunohistochemical images of ten marker genes (AZBA, Kenney *et al*., 2021). Thus, no brain atlas maps gene expression patterns on a large scale. As a promising animal model in behavioral neuroscience, adult zebrafish display more diverse and complex behaviors than larval zebrafish, including associative learning, perceptual decision-making, social behavior, and reproductive behavior (Norton and Bally-Cuif, 2010; Oliveira RF, 2013; Yabuki *et al*., 2016; Chou *et a*l., 2016; Stednitz *et al*., 2018, Okamoto *et al*., 2021). Furthermore, as animal models for various neurological and psychiatric diseases, zebrafish have been actively used in the field of translational research in recent years (Kalueff *et al*., 2014; Stewart *et al*., 2014; Fonseka *et al*., 2016; Fontana *et al*., 2021; Kumar *et al*., 2021). A gene expression atlas of adult zebrafish is needed to expand on these trends.

Here, we report an atlas summarizing the expression patterns of 38 neuropeptide genes and GABAergic/glutamatergic neuronal markers in the adult zebrafish forebrain. Neuropeptides are relatively small peptides that are highly conserved from invertebrates to vertebrates (Holmgren and Jensen, 2001; Elphick *et al*., 2018). They function as neuromodulators and play important roles in various physiological phenomena (Hökfelt *et al*., 2000; van den Pol, 2012; Sohn, 2015; van den Burg and Stoop, 2019). As neuropeptides tend to be expressed in specific types or small localized populations of neurons in the brain, they can also be used as markers for groups of neurons of interest and as genetic tools for visualizing and manipulating these neurons. Indeed, many transgenic zebrafish have been produced using the enhancer/promoter regions of various neuropeptide genes (Liu *et al*., 2003; Levitas-Djerbi *et al*., 2015; Förster *et al*., 2017; Koide *et al*., 2018; Shainer *et al*., 2019; Song *et al*., 2021). By closely examining the expression patterns of neuropeptides and neurotransmitters, we found that some of the hypothalamic nuclei described in the zebrafish reference brain atlas (Wullimann *et al*., 1996) contained multiple subregions with different expression patterns of neuropeptides, which we defined as subnuclei. Furthermore, by comparing the expression patterns of the neuropeptides in the zebrafish hypothalamus with those in the mouse hypothalamus, we identified several nuclei that exhibited similar expression patterns across species.

## 2 Materials and Methods

### 2.1 Animals

Zebrafish (*Danio rerio*) were maintained at 28°C under 14-hour light and 10-hour dark cycle and fed with brine shrimp two to three times a day. Adult wild-type zebrafish (RIKEN-Wako) were used in all the experiments. All experimental procedures were approved by the Animal Care and Use Committee of RIKEN.

### 2.2 cRNA probe synthesis

Coding regions (500–1000 bp) of neuropeptide and neurotransmitter marker genes were PCR-amplified from zebrafish brain cDNA or genomic DNA and cloned into the pGEM-T Easy vector (Promega). Three glutamate decarboxylase genes (*gad1a*, *gad1b*, and *gad2*) were used as marker genes for GABAergic neurons. For the glutamatergic neurons, four vesicular glutamate transporter (*vglut*) genes (*slc17a6a*, *slc17a6b, slc17a7a*, and *slc17a7b*) were used. The genes analyzed and primers used for PCR cloning are listed in Table 1.

**Table 1.**
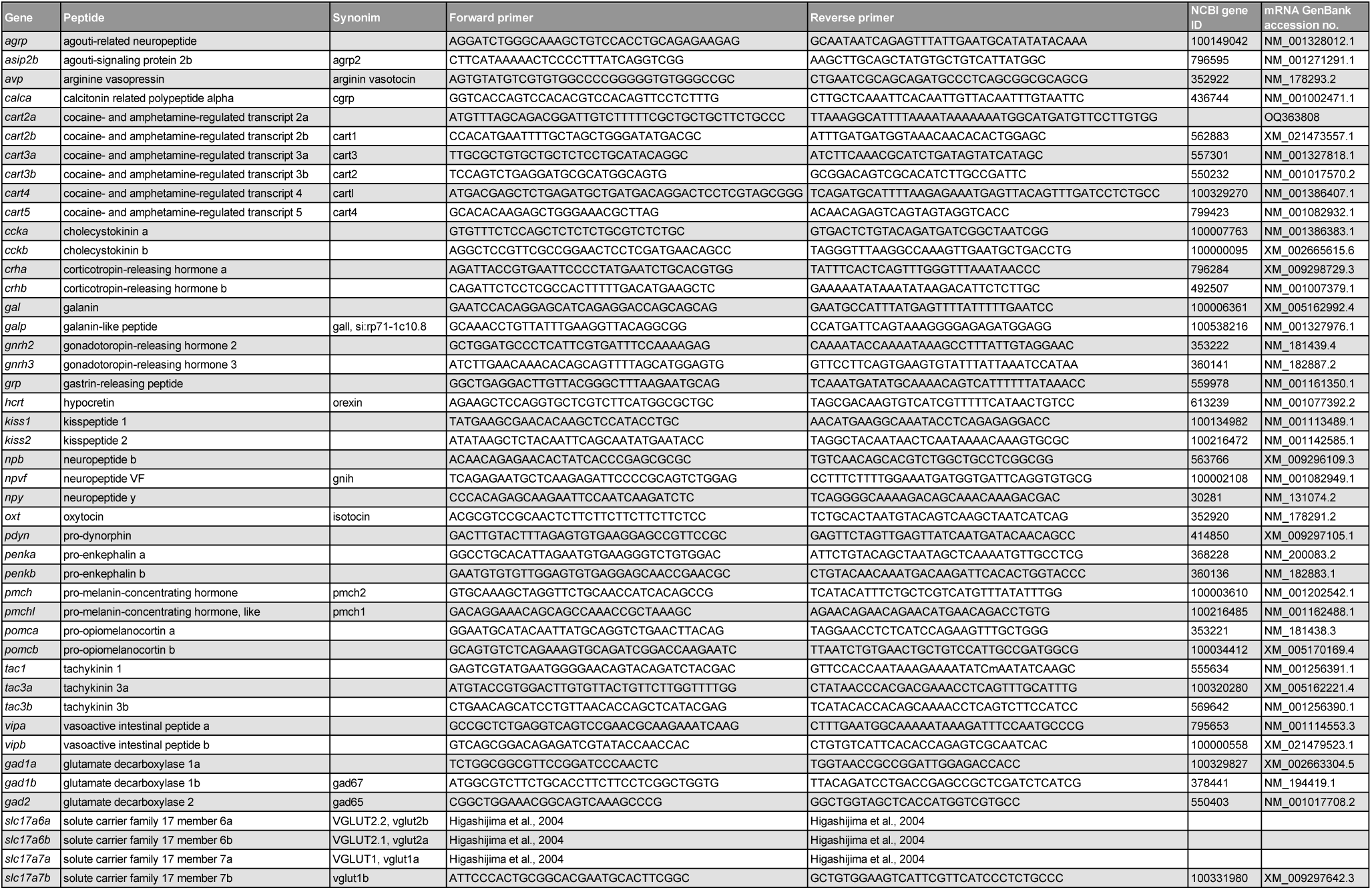
Genes examined in this study and primers used for *in situ* hybridization (ISH) probe synthesis.

The names of the individual genes followed the ZFIN nomenclature, with a few exceptions, for which the correspondences to previous designations are listed in Table 1. One of the *cart* genes was missing from the ZFIN database, and the ZFIN nomenclature did not match the results of the phylogenetic analysis. Therefore, we followed the nomenclature of Cai *et al*. (2015), in which a detailed phylogenetic analysis of *cart* genes was conducted. Zebrafish harbor two galanin genes (*gal* and *gall*); however, the molecular evolutionary positioning of the second galanin-like gene (*gall*, also called *si:rp71-1c10.8*) remains unclear. Hence, we carried out phylogenetic and syntenic analyses of *gall* and revealed that the second galanin-like gene in zebrafish and other teleost species was not a teleost-specific paralogous gene for *gal*, but an orthologous gene for *Galp* in tetrapods (Supplemental Fig. 1). Therefore, it was designated *galp*.

Digoxigenin (DIG)-labeled antisense cRNA probes were synthesized using T7, SP6, or T3 RNA polymerases (Promega) and purified through Micro Bio-Spin Columns with Bio-Gel (BIO-RAD). Plasmids containing *slc17a6a*, *slc17a6b*, and *slc17a7a* genes were kindly provided by Dr. Higashijima (Higashijima *et al*., 2004).

### 2.3 *In situ* hybridization

Brains of adult male zebrafish (3–12 months old) were sampled during the light period and fixed in 4% paraformaldehyde (PFA) in phosphate-buffered saline (PBS) at 4°C overnight. The brains were then incubated in 50%, 70%, 100% ethanol, and Lemosol (Wako) for dehydration, embedded in paraffin, and sectioned into 10 μm-thick serial coronal sections. The sections were deparaffinized in Lemosol and 100%, 90%, 70%, and 50% ethanol. After washing in PBS, the sections were treated using 1 μg/mL proteinase K (Invitrogen) in PBS at 37°C for 15 min and re-fixed in 4% PFA in PBS for 10 min at room temperature (RT). The sections were acetylated, followed by incubation in pre-hybridization buffer [50% formamide, 5× saline sodium citrate (SSC), 50 μg/mL heparin, 0.1% Tween] at 60°C for 1 h. The sections were then hybridized using DIG-labeled neuropeptide gene probes (0.01 ng/μL for the *oxt* gene, 0.5 ng/μL for the rest of the genes) in hybridization buffer [100 μg/μL RNA from yeast (Roche) added to pre-hybridization buffer] at 60°C overnight. The *gad* mix (a mixture of *gad1a*, *gad1b*, and *gad2* probes; 0.5 ng/μL each) was used as a marker for GABAergic neurons. Moreover, *vglut* mix (a mixture of *slc17a6a*, *slc17a6b, slc17a7a*, and *slc17a7b* probes; 0.5 ng/μL each) was used as a marker for glutamatergic neurons. The sections were then washed in 5× SSC buffer and 0.2× SSC at 60°C. After washing in 0.2× SSC and maleic acid buffer (MAB; 0.1 M maleic acid, 0.15 M NaCl, 0.19 M NaOH), the sections were blocked with 2% blocking reagent (Roche) and 5% normal goat serum in MAB for 30 min at RT and incubated with alkaline phosphatase-conjugated anti-DIG antibody (1:10000 for *oxt*, 1:1000 for other genes, Sigma-Aldrich) at 4°C overnight. For color development, the sections were washed in MAB and color development buffer (0.1 M Tris-HCl pH 9.5, 0.05 M MgCl_2_, 0.01 M NaCl) and then incubated with 5-bromo-4-chloro-3-indolyl phosphate/nitro blue tetrazolium (BCIP/NBT) solution at 30°C for 0.5–7 h. The color development was stopped by washing with PBS and water. Some sections (at least one individual fish for each gene) were counterstained using 1% methyl green stain solution (Muto Pure Chemicals) for 3–5 min and washed in water. The sections were then mounted using Aqua-Poly/Mount (Polysciences) or Immu-mount (Thermo Fisher Scientific).

### 2.4 Image acquisition

Whole images of brain sections on glass slides were captured using a NanoZoomer HT slide scanner (Hamamatsu Photonics). The pictures were cut out for each brain plane using NDP.view2 (Hamamatsu Photonics) and trimmed using Adobe Photoshop CS6 (Adobe Inc.).

### 2.5 Database construction

The Adult Zebrafish Brain Gene Expression Database (https://ssbd.riken.jp/azebex/) was constructed based on the SSBD (Systems Science of Biological Dynamics) platform (Tohsato *et al*., 2016). SSBD:repository was used as the database infrastructure. OMERO (Open Microscopy Environment) system (Allan *et al*., 2012) was used to preview the images.

### 2.6 Phylogenetic and syntenic analyses

Phylogenetic and syntenic analyses were performed to examine homologous relationships between the two galanin-like genes in various animal species. The GenBank accession numbers of the genes used for phylogenetic analysis are summarized in Supplemental Table 1. The amino acid sequences of the galanin-like genes in each species were aligned using ClustalW, and a phylogenetic tree was constructed using the maximum likelihood method. The analysis was performed using Molecular Evolutionary Genetics Analysis (MEGA11) software (version 11) (Tamura *et al*., 2021). For syntenic analysis, genomic information for each species was obtained from the Ensembl or NCBI databases.

### 2.7 Comparison of the hypothalamic nuclei between zebrafish and mouse

To examine the similarity in the hypothalamic nuclei between zebrafish and mice, we compiled a table of gene expression in each nucleus (expressed = 1, not expressed = 0). Comparisons were made for genes that were expressed in both the zebrafish and mouse hypothalamus. Expression profiles of neuropeptide genes in the mouse hypothalamus were obtained from the Allen Brain Atlas (Lein *et al*., 2007) and from previous studies (*Agrp*, Deem *et al*., 2021; *Avp*, Smith *et al*., 1996; Otero-García *et al*., 2015; *Crh*, Alon *et al*., 2009; Keegan *et al*., 1994; *Galp*, Juréus *et al*., 2001; *Oxt*, Otero-García *et al*., 2015; *Pmch*, Croizier *et al*., 2010; Diniz *et al*., 2019; *Npb*, Tanaka *et al*., 2003; *Hcrt*, Peyron *et al*., 1998; *Pomc*, Williams *et al*., 2010; Cowley *et al*., 2001; *Tac1*, Navarro *et al*., 2014; *Tac2*, Duarte *et al*., 2005). The expression of the zebrafish *gad* mix was compared to that of *Gad1* and *Gad2* in mice. Zebrafish *vglut* mix expression was compared with *Vglut2* expression in mice because *Vglut1* was not expressed in the mouse hypothalamus. Because *gad*/*Gad* and *vglut*/*Vglut* are more or less expressed in almost all regions, the expression patterns of *gad* and *vglut* were compared, and those with a higher number of expressing neurons were counted as positive. Both were considered positive when they were equally expressed. When neither was expressed, both were considered negative. The nomenclature of the mouse hypothalamic nuclei follows that of Paxinos and Franklin (2019). Zebrafish often have multiple homologous genes corresponding to a single mouse gene, mainly because of teleost-specific genome duplication (Jaillon *et al*., 2004, Kassahn *et al*., 2009). Teleost-specific paralogous genes expressions, that have been shown to have arisen from teleost-specific genome duplication, were added together (*ccka*/*cckb*, Dupré and Tostivint, 2014; Kurokawa *et al*., 2003; *crha*/*crhb*, Grone and Maruska, 2015; *penka*/*penkb*, Bojnik *et al*., 2011; *pomca*/*pomcb*, Forlano and Cone, 2007; *tac3a*/*tac3b*, Zhou *et al*., 2012; *vipa*/*vipb*, Nam *et al*., 2009). The *cart* genes were expressed in both zebrafish and mice but were not included in the comparison analysis because none of the *cart* genes found in the present zebrafish genome data were considered homologous to the mouse *Cart* gene (Cai *et al*., 2015). Furthermore, zebrafish and mouse hypothalamic nuclei and neuropeptide/neurotransmitter genes were clustered using Ward’s method based on Euclidean distance using the hclust function in R: A Language and Environment for Statistical Computing (R Foundation for Statistical Computing).

## 3. Results

### 3.1 Neuropeptide expressions in the adult zebrafish forebrain

Expression patterns of 38 neuropeptide genes, *agrp*, *asip2b*, *avp*, *calca*, *cart2a*, *cart2b*, *cart3a*, *cart3b*, *cart4*, *cart5*, *ccka*, *cckb*, *crha*, *crhb*, *gal*, *galp*, *gnrh2*, *gnrh3*, *grp*, *kiss1*, *kiss2*, *npb*, *npvf*, *npy*, *hcrt*, *oxt*, *pdyn*, *penka*, *penkb*, *pmch*, *pmchl*, *pomca*, *pomcb*, *tac1*, *tac3a*, *tac3b*, *vipa*, *vipb*, glutamatergic neuron marker gene mix (*slc17a6a*, *slc17a6b*, *slc17a7a*, and *slc17a7b*), and GABAergic neuron marker gene mix (*gad1a*, *gad1b*, and *gad2*) were examined by *in situ* hybridization throughout the adult zebrafish forebrain (telencephalon and diencephalon). Fig. 1 shows a schematic of the zebrafish brain and the location of each plane. Figs. 2–20 show schematics of the coronal planes and microscopic images of the genes that are expressed in each plane. Schematics were based on the zebrafish brain atlas (Wullimann *et al*., 1996) with several modifications. The division of nuclei in the dorsal telencephalic area was modified according to Ganz *et al*. (2014) and Yáñez *et al*. (2022) (Figs. 3–10). The intermediate nucleus of the ventral telencephalon (Vi) was added, as described by Biechl *et al*. (2017) (Figs. 9, 10). Subregions in the dorsal habenula (Had), medial Had (Hadm), and lateral Had (Hadl) were added according to Agetsuma *et al*. (2010) (Figs. 11–13). The nucleus lateralis tuberis (NLT) in the hypothalamus was added as in recent papers in zebrafish (Akash *et al*., 2014) or other teleost species (Karoubi *et al*., 2016) (Figs. 14, 15). The intercalated nucleus (IC) was added, as described by Mueller and Guo (2009). The intermediate nucleus (IN) was added, and the positions of the paraventricular organ (PVO) and posterior tuberal nucleus (PTN) were altered according to Rink and Wullimann (2001) (Figs. 15, 16). The neuropeptide expression profiles in each nucleus are summarized in Tables 2–6. All scanned data of the whole glass slides are available in the Adult Zebrafish Brain Gene Expression Database (https://ssbd.riken.jp/azebex). The database allows users to view whole-brain scan data online. Raw data in .ndpi format can also be downloaded from the database and viewed using NDP.view2, which is free software. The database also provides images of sections in the hindbrain and expression data in female zebrafish.

**Figure 1.** Schematic of the zebrafish brain viewed from the lateral side. The anterior is to the left, and the dorsal is to the top. Lines 1–19 indicate the locations of each plane in Figs. 2–20.

**Figure 2.**
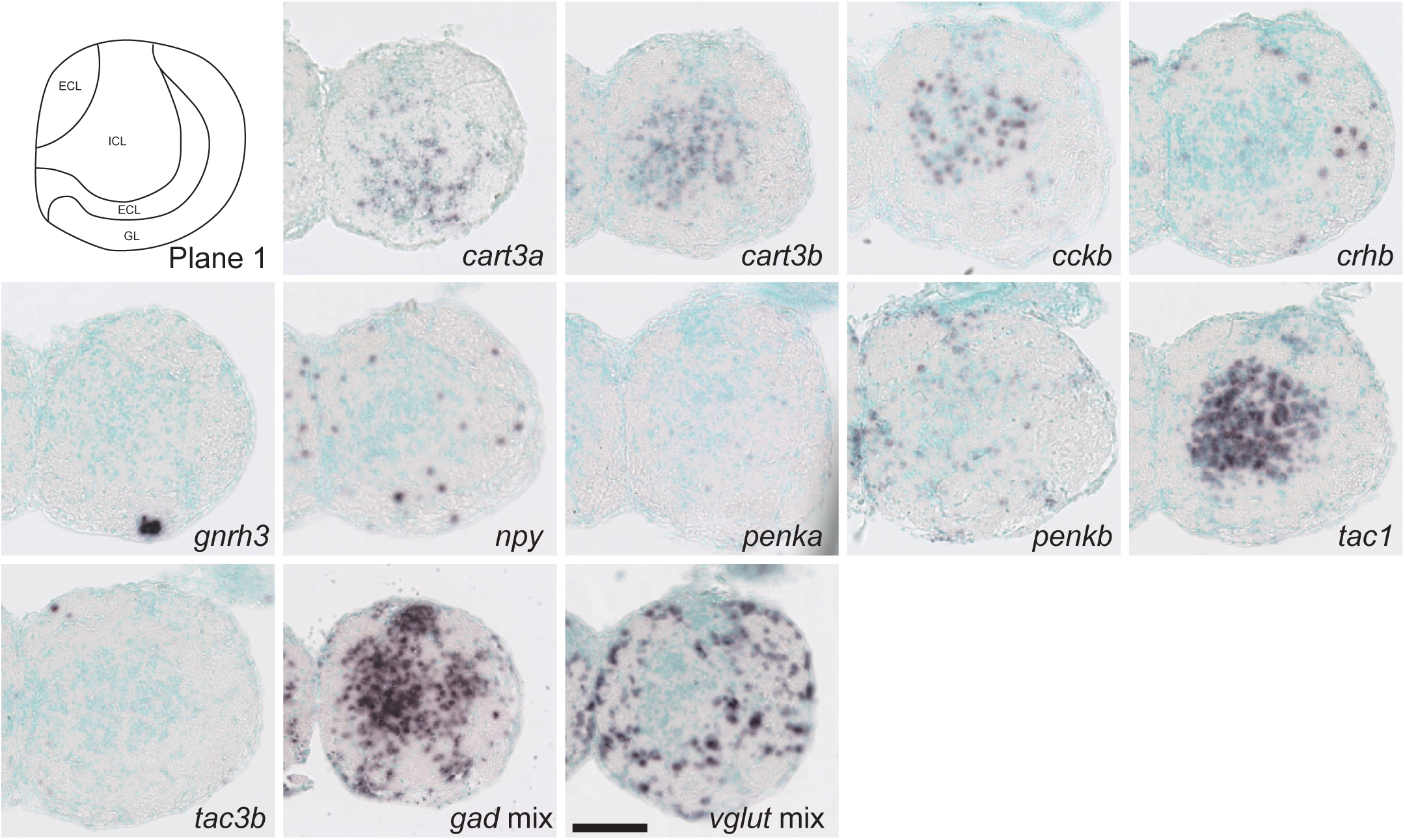
Schematic of plane 1 and photographs of ISH results of genes expressed in plane 1. Scale bar = 100 μm.

**Figure 3.**
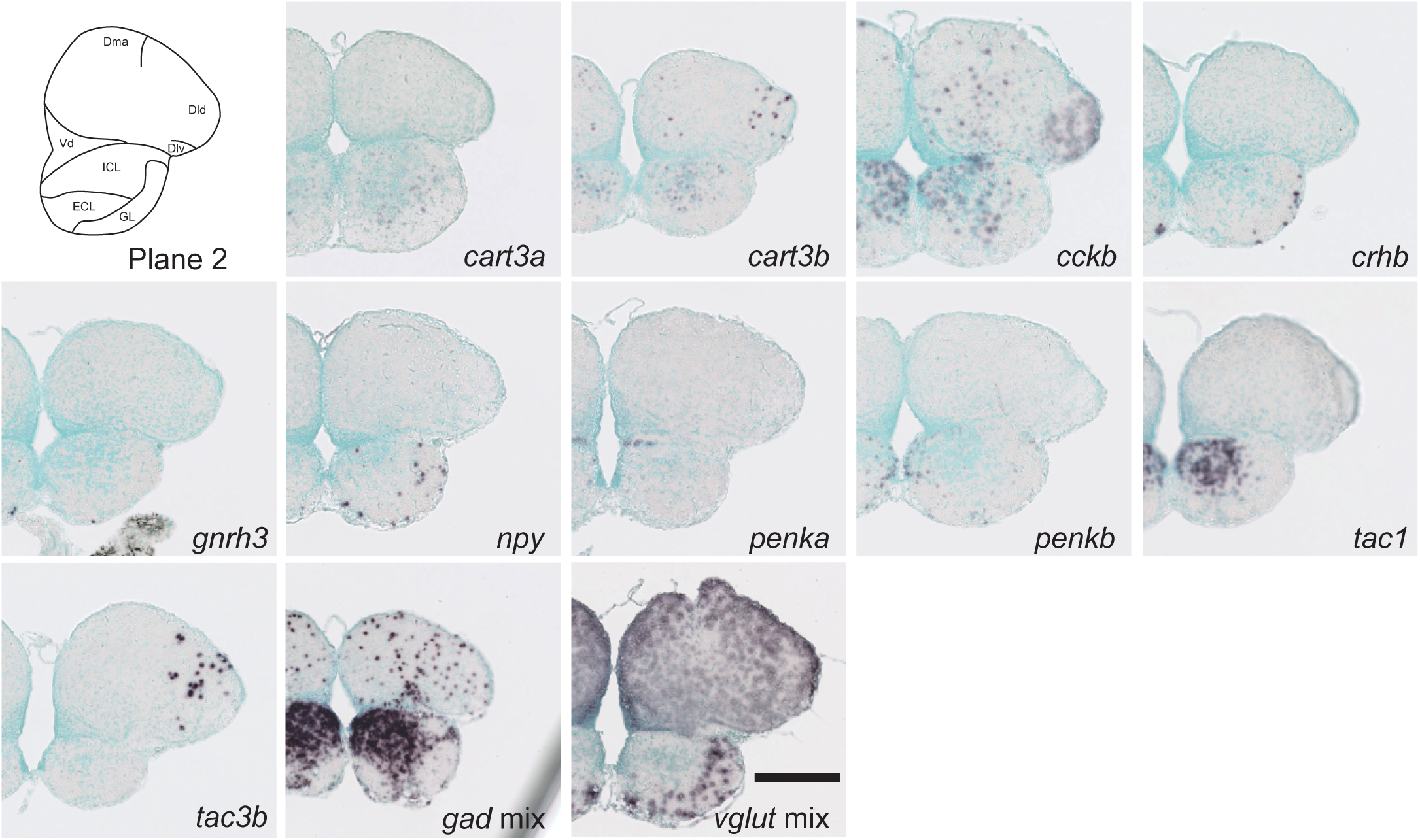
Schematic of plane 2 and photographs of ISH results of genes expressed in plane 2. Scale bar = 200 μm.

**Figure 4.**
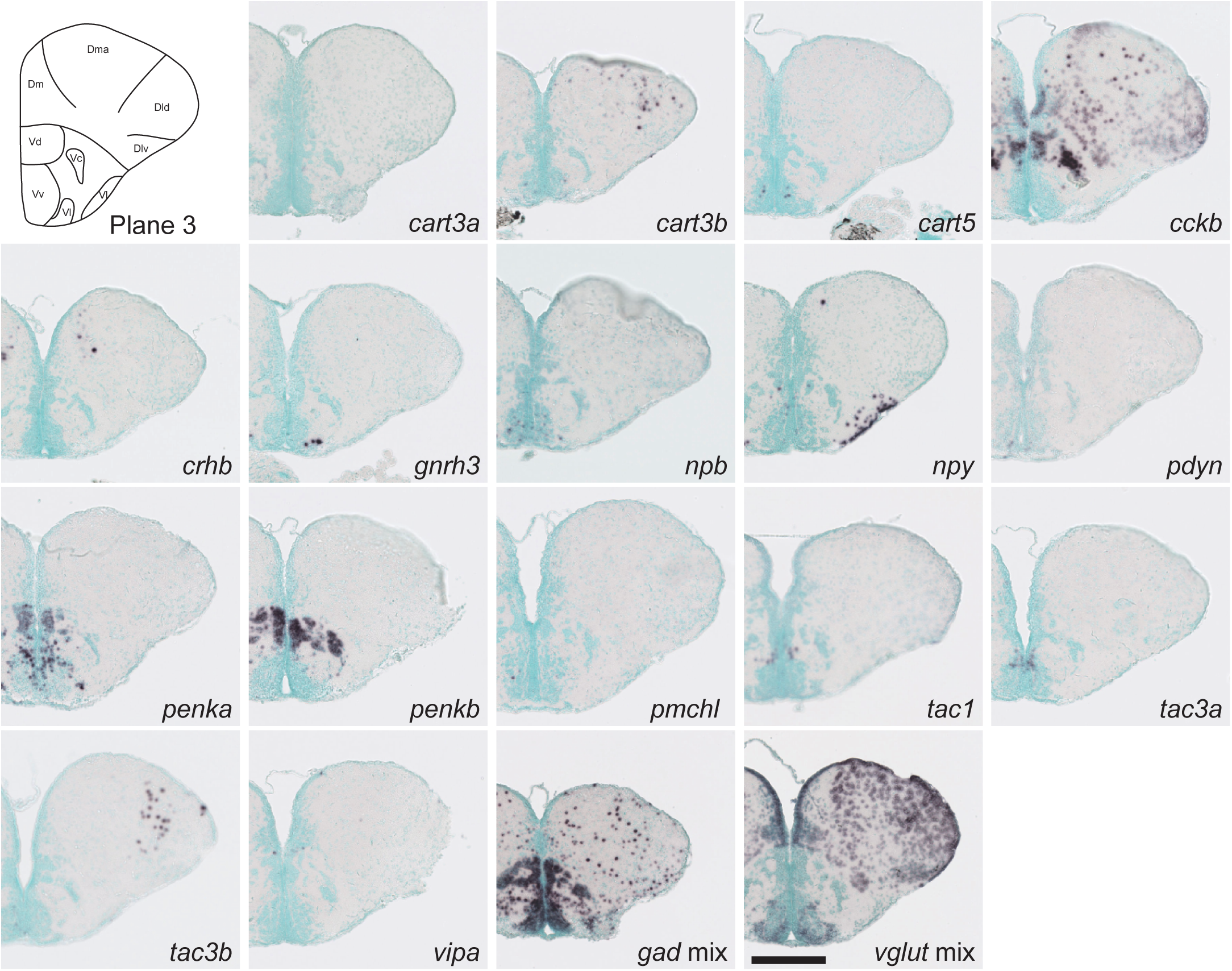
Schematic of plane 3 and photographs of ISH results of genes expressed in plane 3. Scale bar = 200 μm.

**Figure 5.**
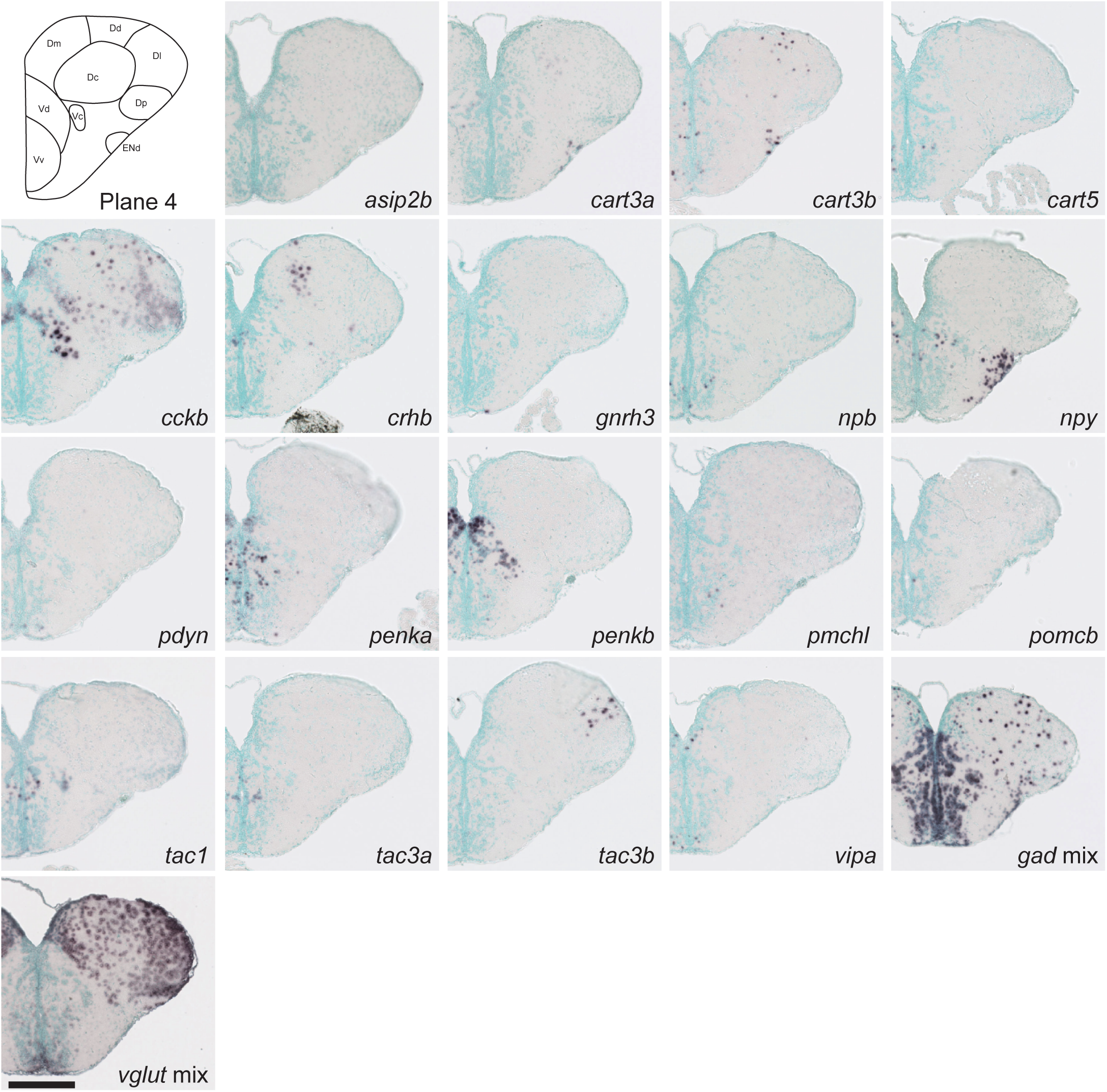
Schematic of plane 4 and photographs of ISH results of genes expressed in plane 4. Scale bar = 200 μm.

**Figure 6.**
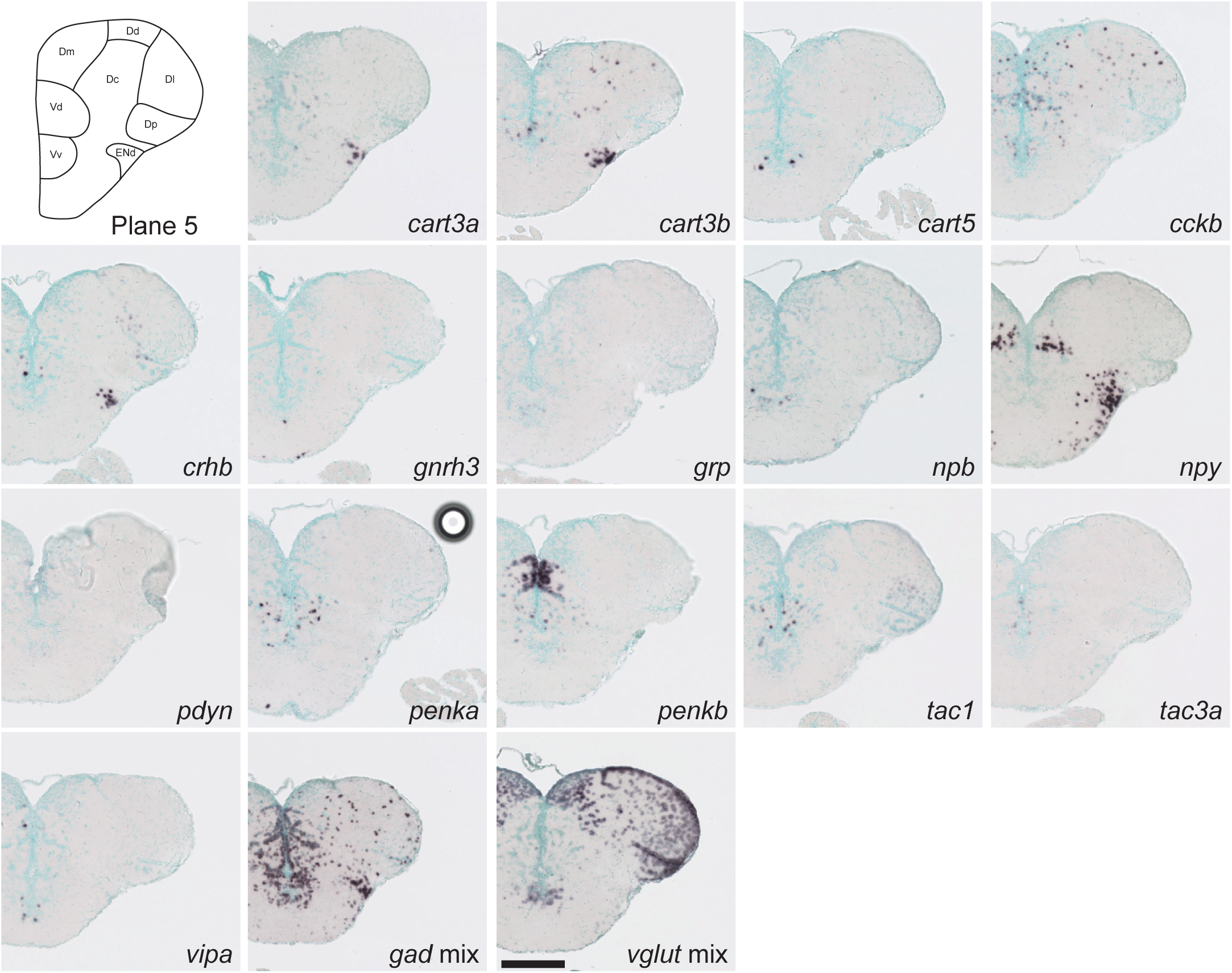
Schematic of plane 5 and photographs of ISH results of genes expressed in plane 5. Scale bar = 200 μm.

**Figure 7.**
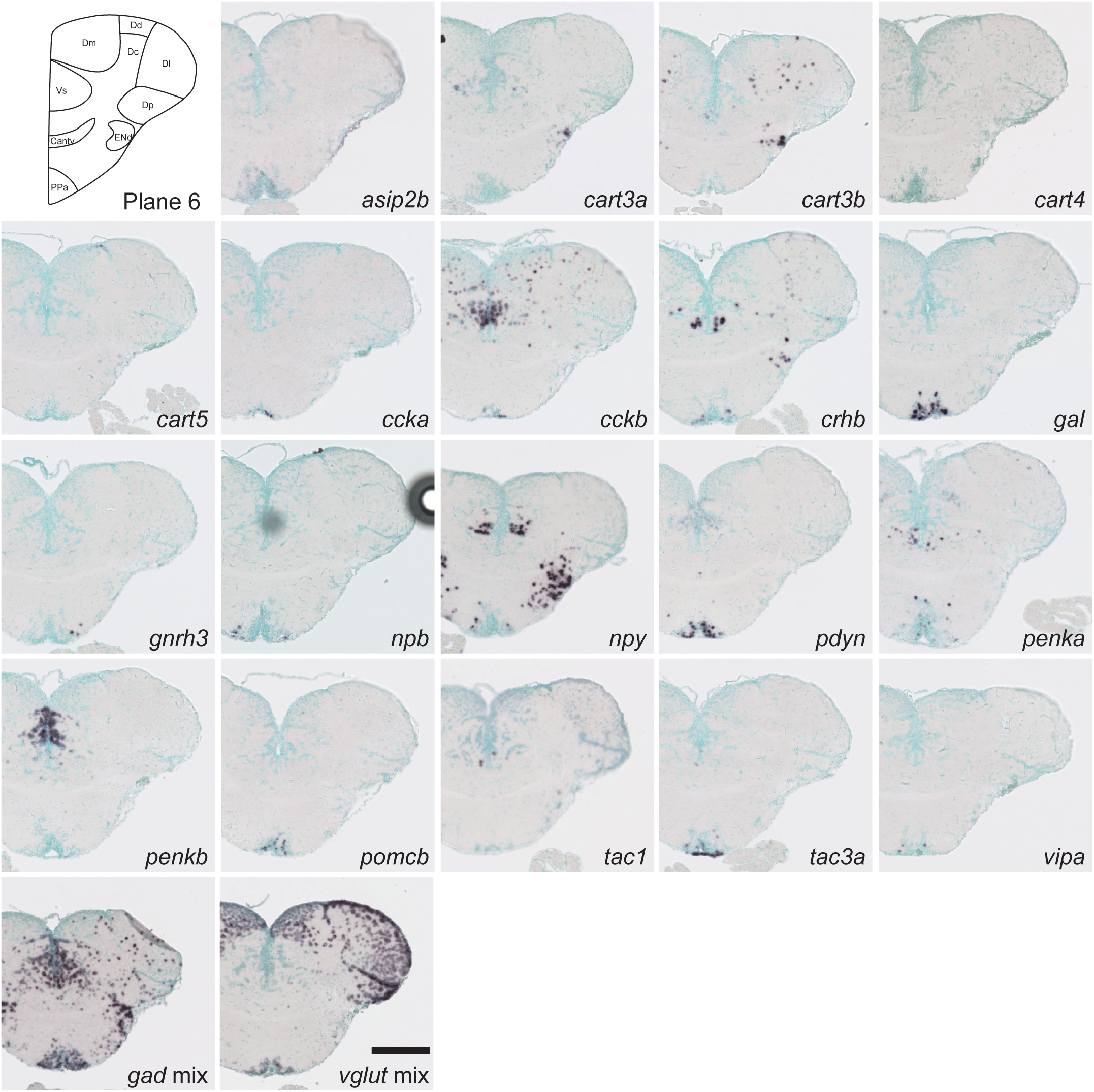
Schematic of plane 6 and photographs of ISH results of genes expressed in plane 6. Scale bar = 200 μm.

**Figure 8.**
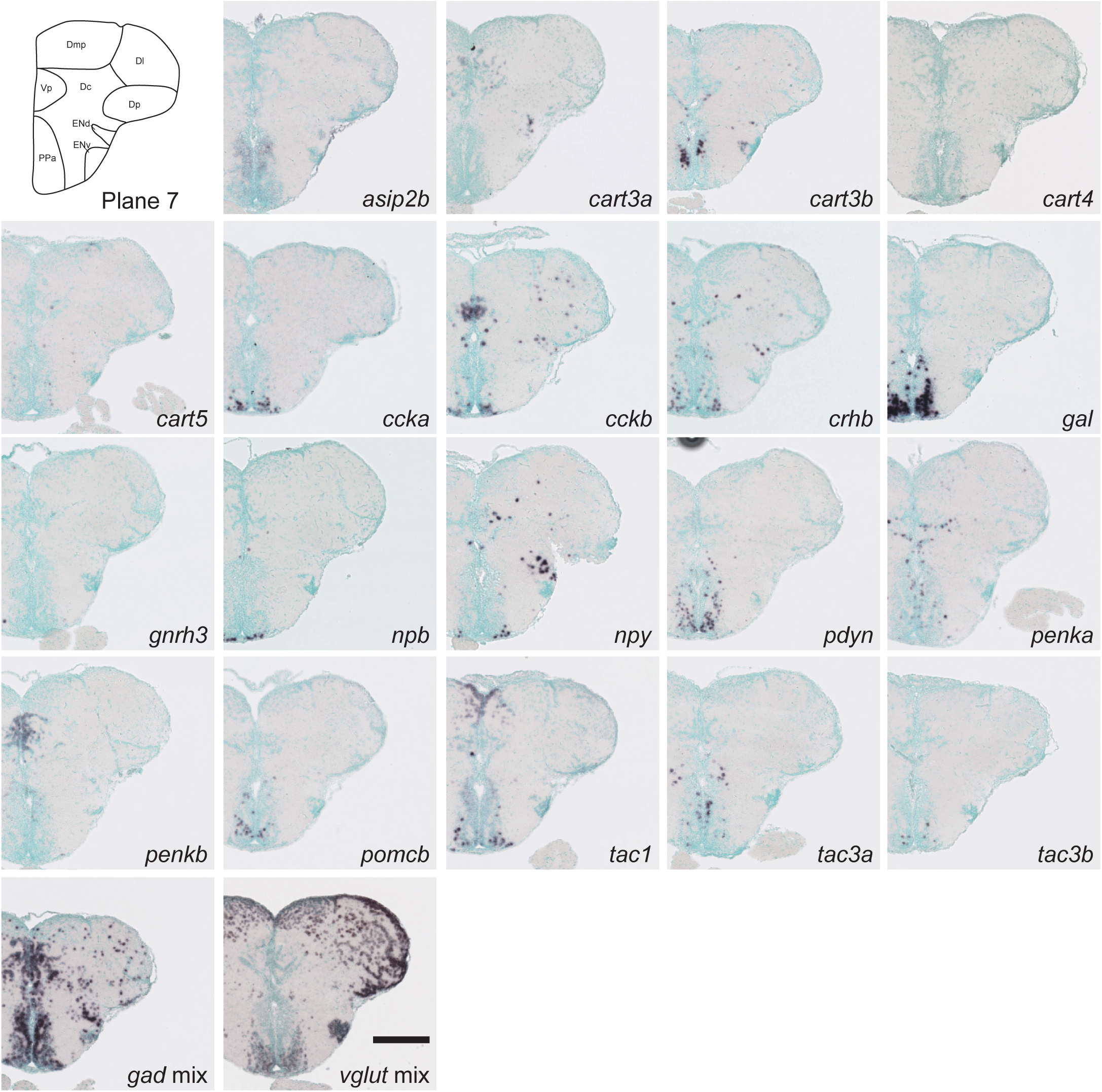
Schematic of plane 7 and photographs of ISH results of genes expressed in plane 7. Scale bar = 200 μm.

**Figure 9.**
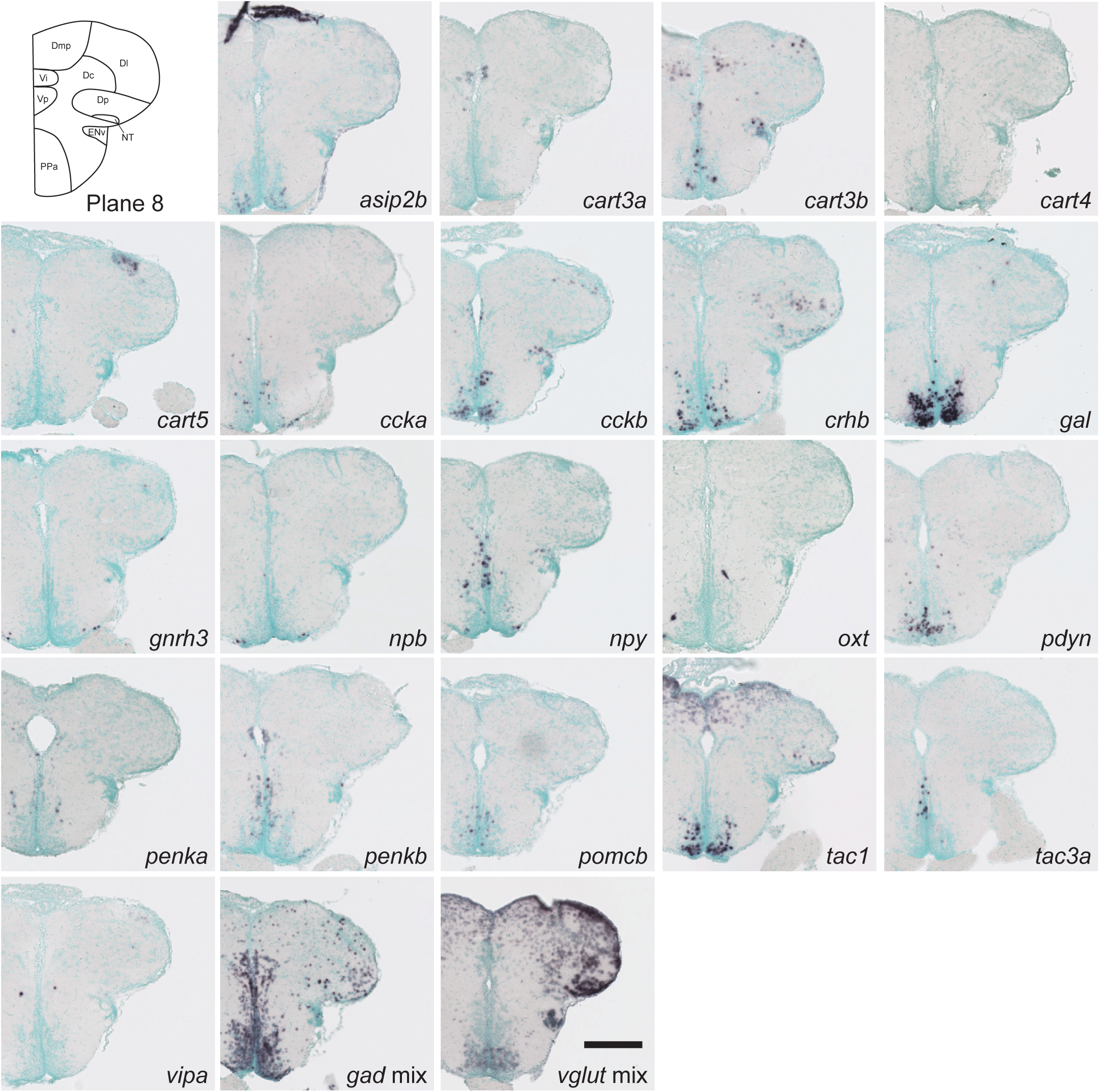
Schematic of plane 8 and photographs of ISH results of genes expressed in plane 8. Scale bar = 200 μm.

**Figure 10.**
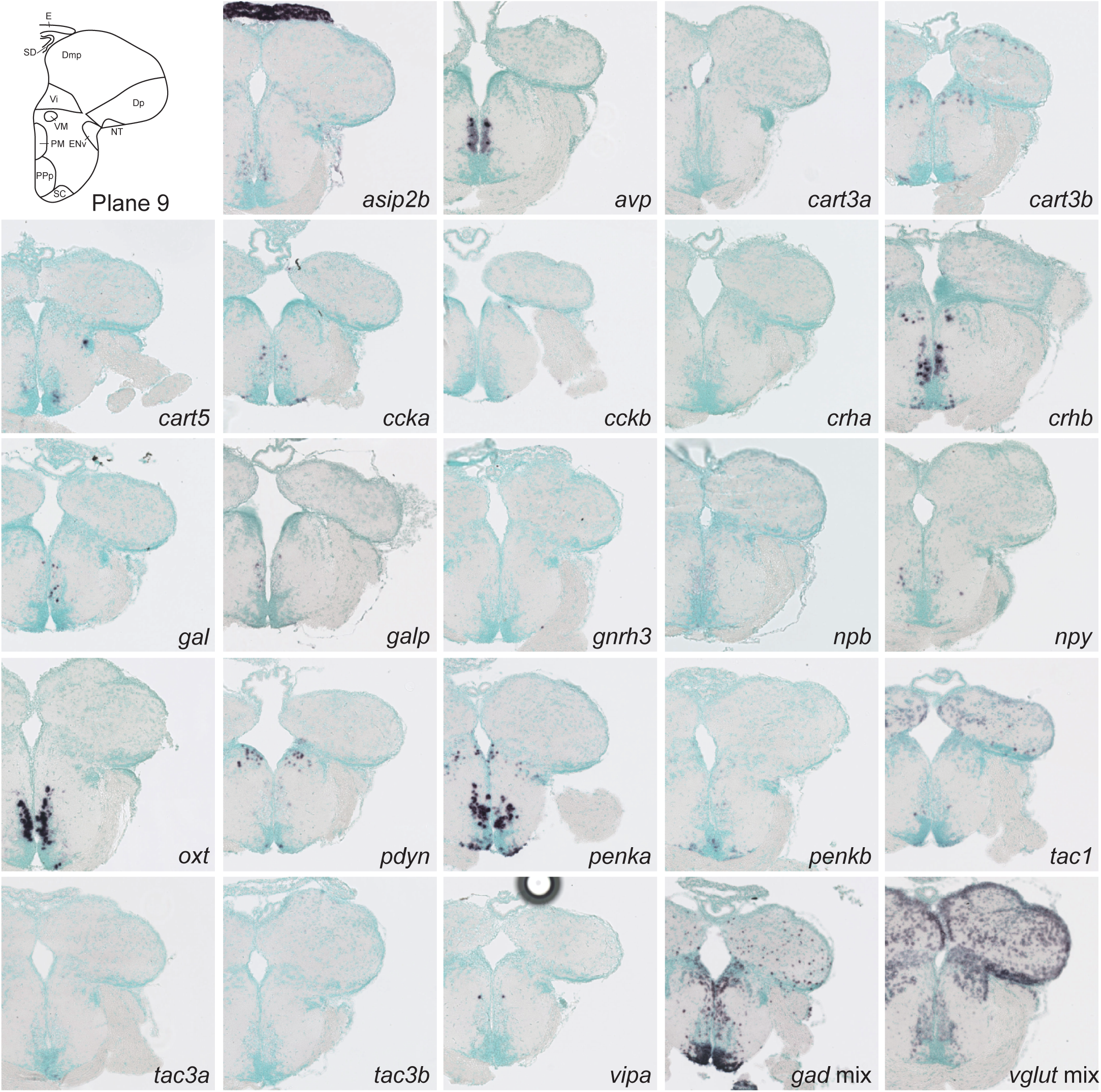
Schematic of plane 9 and photographs of ISH results of genes expressed in plane 9. Scale bar = 200 μm

**Figure 11.**
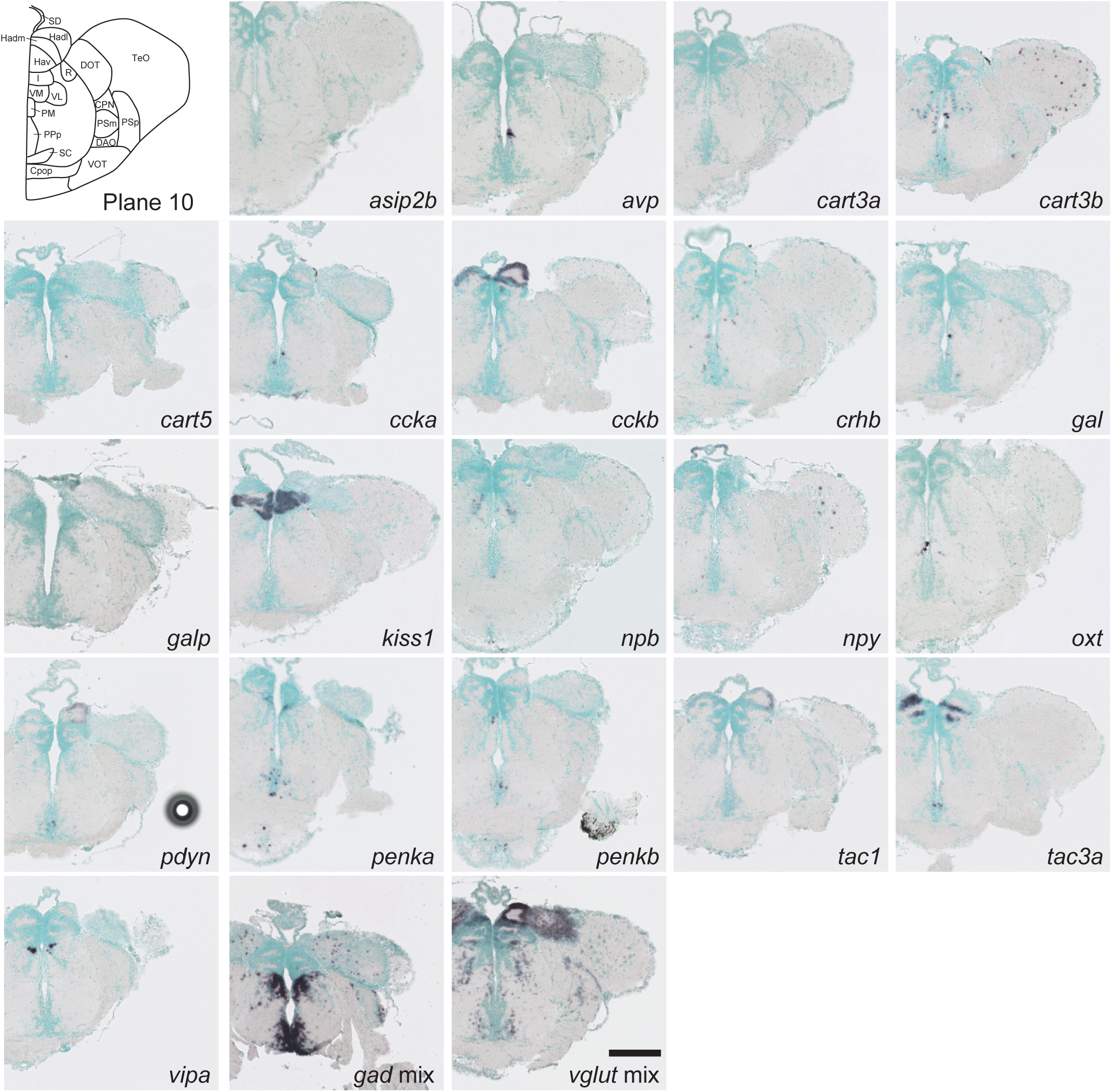
Schematic of plane 10 and photographs of ISH results of genes expressed in plane 10. Scale bar = 200 μm.

**Figure 12.**
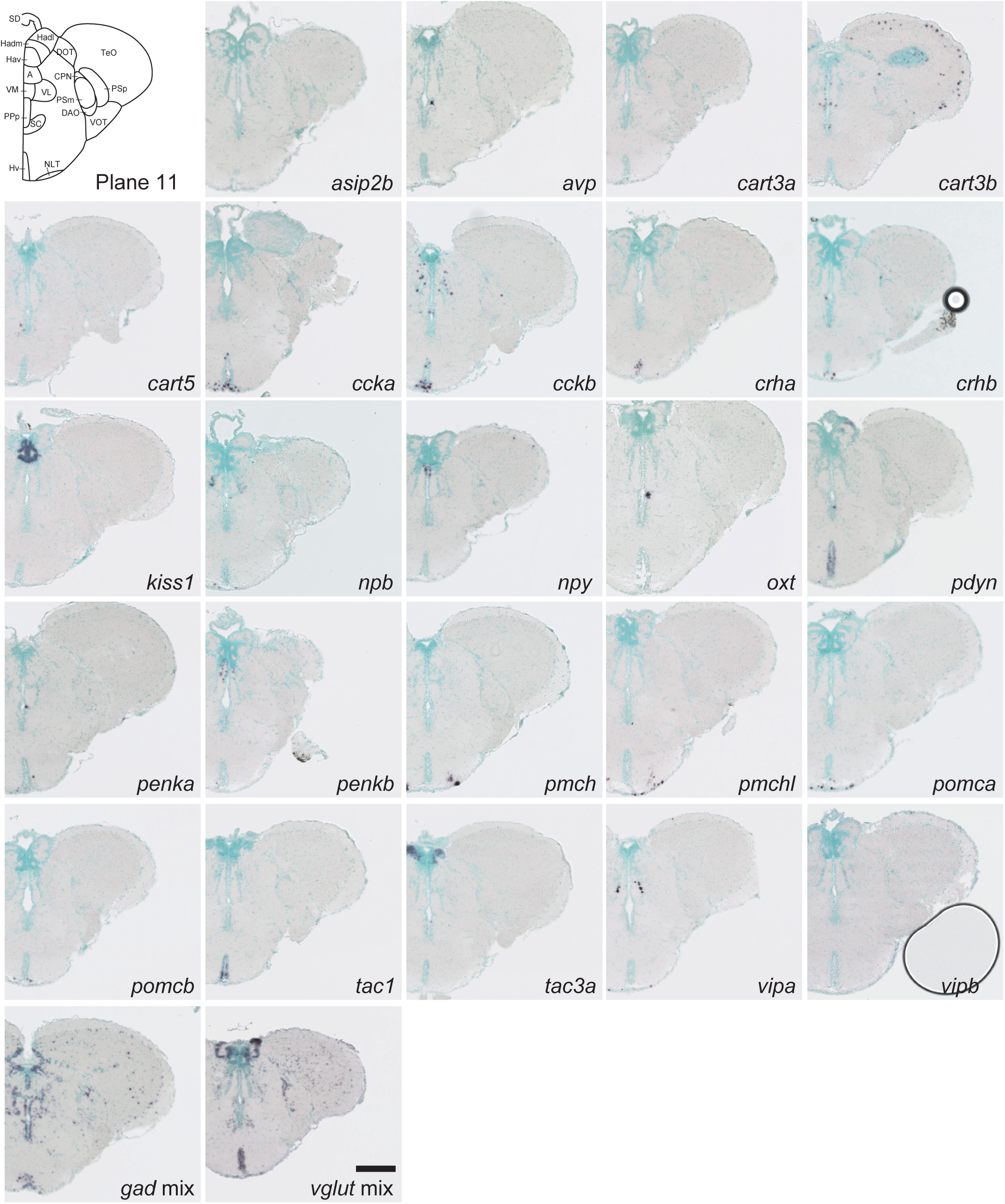
Schematic of plane 11 and photographs of ISH results of genes expressed in plane 11. Scale bar = 200 μm.

**Figure 13.**
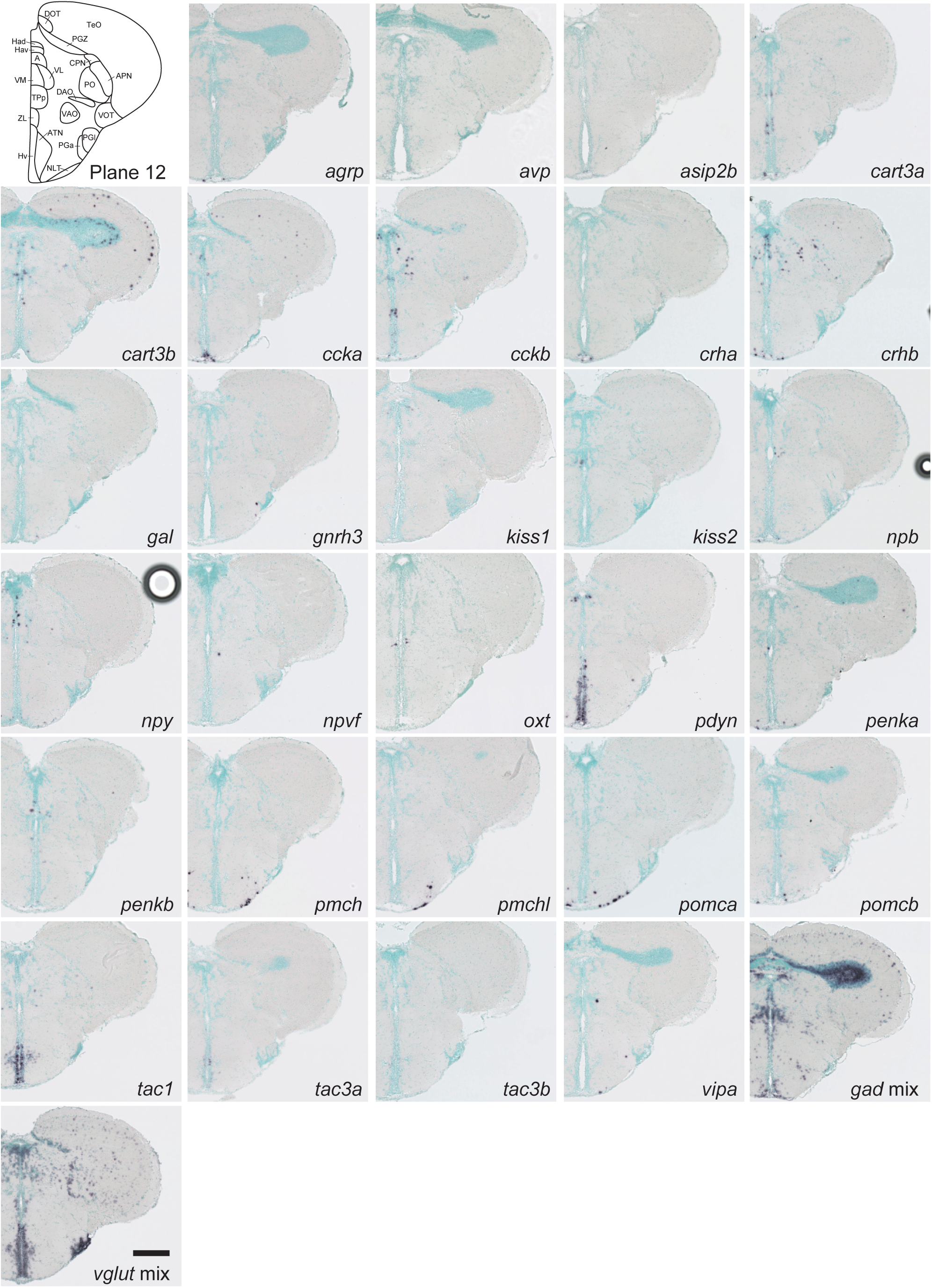
Schematic of plane 12 and photographs of ISH results of genes expressed in plane 12. Scale bar = 200 μm.

**Figure 14.**
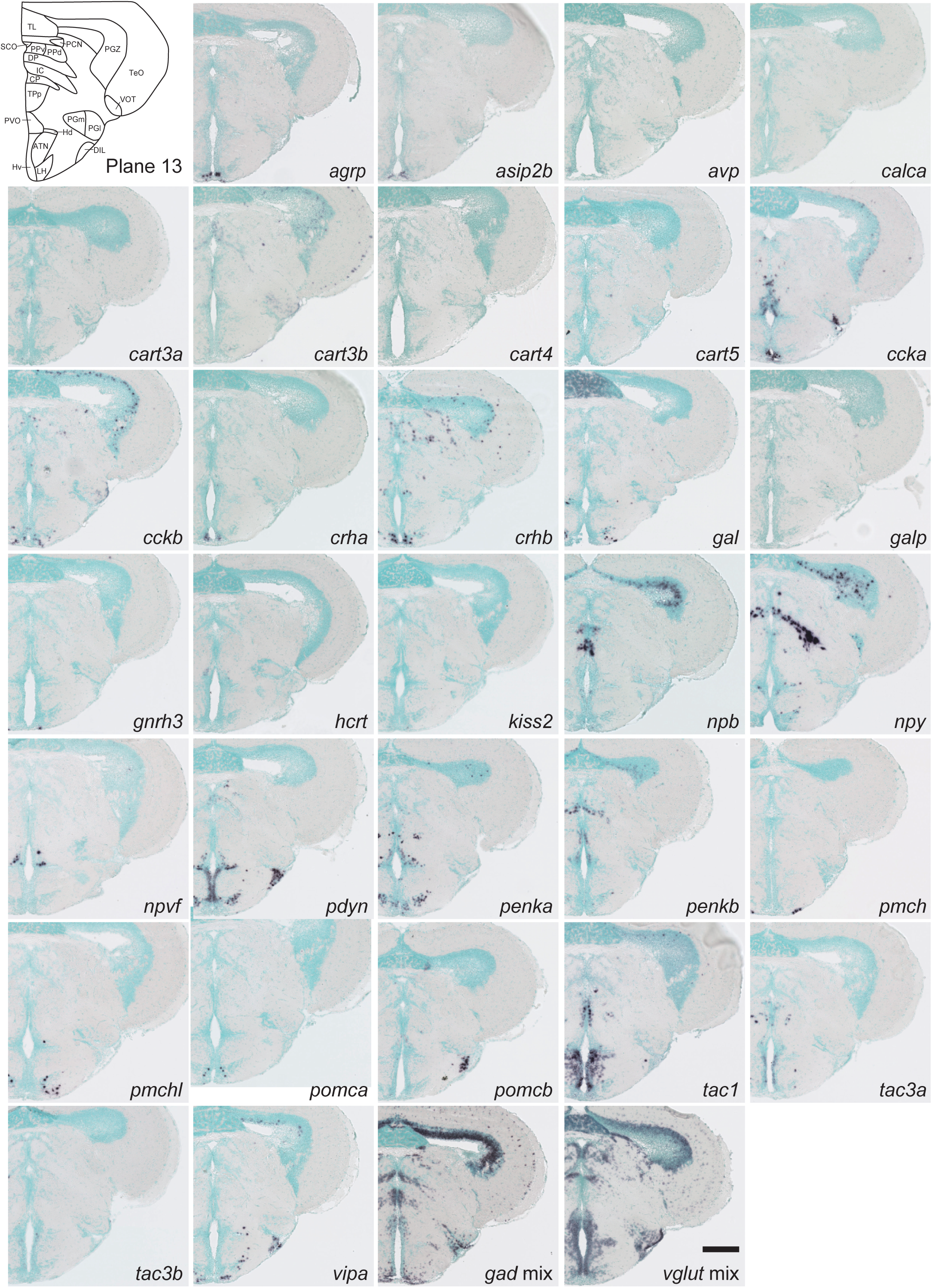
Schematic of plane 13 and photographs of ISH results of genes expressed in plane 13. Scale bar = 200 μm.

**Figure 15.**
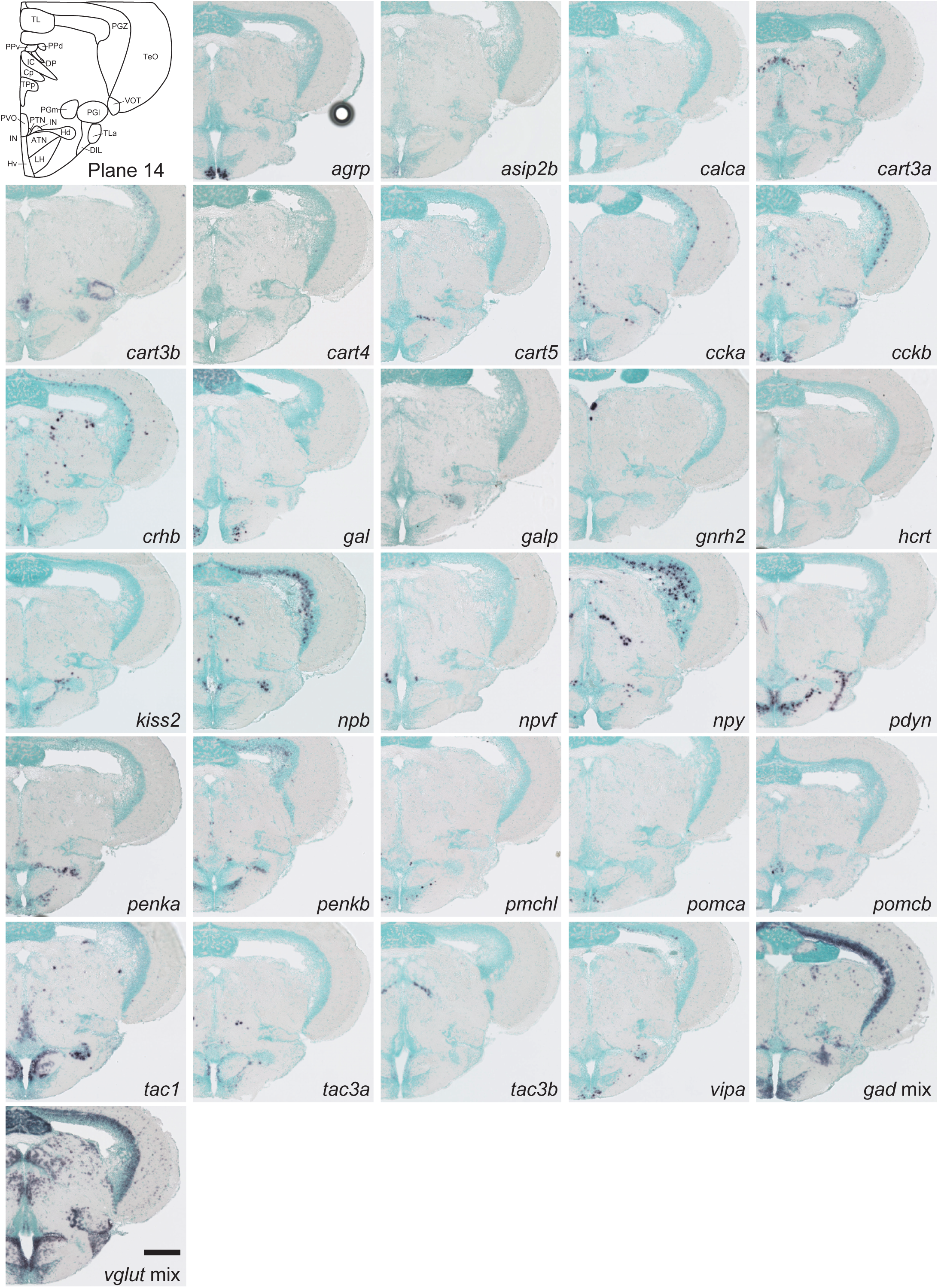
Schematic of plane 14 and photographs of ISH results of genes expressed in plane 14. Scale bar = 200 μm.

**Figure 16.**
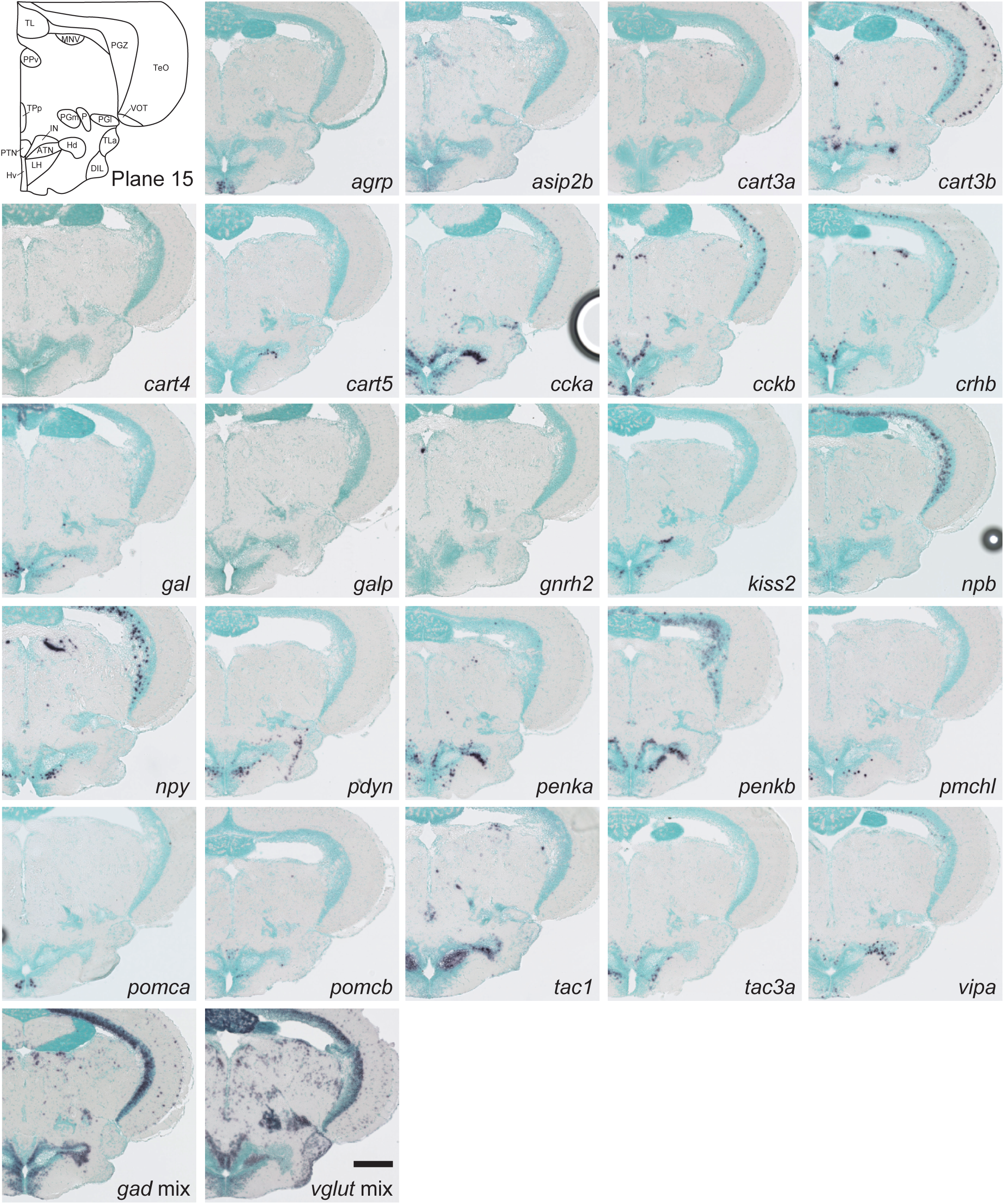
Schematic of plane 15 and photographs of ISH results of genes expressed in plane 15. Scale bar = 200 μm.

**Table 2.** The summary of gene expressions in the olfactory bulb and dorsal telencephalon.

|  | Olfactory bulb |  |  | Dorsal telencephalon |  |  |  |  |  |  |  |  |
| --- | --- | --- | --- | --- | --- | --- | --- | --- | --- | --- | --- | --- |
|  | ICL | ECL/GL | Terminal<br>nerve<br>neruon | Dma | Dld | Dlv | Dm | Dd | Dc | Dp | DI | Dmp |
| agrp | - | - | - | - | - | - | - | - | - | - | - | - |
| asip2b | - | - | - | - | - | - | - | - | - | - | - | - |
| avp | - | - | - | - | - | - | - | - | - | - | - | - |
| calca | - | - | - | - | - | - | - | - | - | - | - | - |
| cart2a | - | - | - | - | - | - | - | - | - | - | - | - |
| cart2b | - | - | - | - | - | - | - | - | - | - | - | - |
| cart3a | + | + | - | - | - | - | - | - | + | - | - | - |
| cart3b | + | - | - | + | + | - | + | + | + | - | + | + |
| cart4 | - | - | - | - | - | - | - | - | - | - | - | - |
| cart5 | - | - | - | - | - | - | + | - | - | - | + | - |
| ccka | - | - | - | - | - | - | - | - | - | - | - | - |
| cckb | + | - | - | + | + | + | + | + | + | + | + | - |
| crha | - | - | - | - | - | - | - | - | - | - | - | - |
| crhb | + | + | - | - | - | - | + | + | + | + | + | - |
| gal | - | - | - | - | - | - | - | - | - | - | - | - |
| galp | - | - | - | - | - | - | - | - | - | - | - | - |
| gnrh2 | - | - | - | - | - | - | - | - | - | - | - | - |
| gnrh3 | - | - | + | - | - | - | - | - | - | - | - | - |
| grp | - | - | - | - | - | - | - | - | - | - | - | - |
| hcrt | - | - | - | - | - | - | - | - | - | - | - | - |
| kiss1 | - | - | - | - | - | - | - | - | - | - | - | - |
| kiss2 | - | - | - | - | - | - | - | - | - | - | - | - |
| npb | - | - | - | - | - | - | - | - | - | - | - | - |
| npvf | - | - | - | - | - | - | - | - | - | - | - | - |
| npy | - | + | - | - | - | - | - | - | + | - | - | - |
| oxt | - | - | - | - | - | - | - | - | - | - | - | - |
| pdyn | - | - | - | - | - | - | - | - | - | - | - | - |
| penka | + | - | - | - | - | - | + | - | - | - | + | + |
| penkb | + | + | - | - | - | - | - | - | - | - | - | - |
| pmch | - | - | - | - | - | - | - | - | - | - | - | - |
| pmchl | - | - | - | - | - | - | - | - | - | - | - | - |
| pomca | - | - | - | - | - | - | - | - | - | - | - | - |
| pomcb | - | - | - | - | - | - | - | - | - | - | - | - |
| tac1 | + | - | - | - | - | - | - | - | - | + | + | + |
| tac3a | - | - | - | - | - | - | - | - | - | - | - | - |
| tac3b | - | + | - | - | + | - | - | - | - | - | - | - |
| vipa | - | - | - | - | - | - | + | - | - | - | + | - |
| vipb | - | - | - | - | - | - | - | - | - | - | - | - |
| gad mix | + | + | - | + | + | + | + | + | + | + | + | + |
| vglut mix | + | + | - | + | + | + | + | + | + | + | + | + |

#### 3.1.1 Telencephalon

##### 3.1.1.1 Olfactory bulb

In the zebrafish olfactory bulb, the glomerular layer (GL) and the external cellular layer (ECL) contain two types of projection neurons, mitral cells and ruffed cells, and two types of interneurons, periglomerular cells and short-axon cells. The internal cellular layer (ICL) mainly contains GABAergic interneurons and granule cells (Olivares and Schmachtenberg, 2019). The distribution of GABAergic and glutamatergic neurons was consistent with that of these distinct cell types (Figs. 2 and 3, Table 2). *crhb* was strongly expressed in a small number of large-diameter cells in ECL/GL, which presumably are a subset of mitral cells, as was reported in mammals (Bassett *et al*., 1992) (Figs. 2, 3, Table 2). *penkb* was expressed in small-diameter cells surrounding glomeruli in GL/ECL and ICL, which are likely periglomerular cells and granule cells, respectively, as is the case in mammals (Matsutani *et al*., 1989) (Figs. 2, 3, Table 2). *npy*-positive cells in GL/ECL might be short-axon cells, as described in mammals (Scott *et al*., 1987). *cart3b*, *cckb*, and *tac1* were expressed intensely and exclusively in ICL cells, which are likely granule cells. *gnrh3* was expressed in terminal nerve neurons, with a very small number of cells located in the ventral-most region of the olfactory bulb (Figs. 2 and 3, Table 2).

##### 3.1.1.2 Dorsal telencephalon

The dorsal telencephalic area (D) contains nine nuclei, including several newly defined nuclei (Ganz *et al*., 2014; Yáñez *et al*., 2022) (Figs. 3–10, Table 2). All nuclei showed distinct neuropeptide expression patterns, some of which validated the new definition of nuclei (Figs. 3–10, Table 2). For example, *cart3b* and *tac3b* were selectively expressed in the dorsal part of the lateral zone of D (Dld), but not in the ventral part of the lateral zone of D (Dlv) (Figs. 3 and 4, Table 2). *tac1* was strongly expressed in the posterior part of the medial zone of D (Dmp) but not in the medial zone of D (Dm) (Figs. 4–10, Table 2). Dmp was also characterized by the absence of *cckb* expression (Fig. 8–10, Table 2). The dorsal telencephalon was characterized by high *vglut* expression and low *gad* expression (Figs. 3–10). The dorsal telencephalon expressed fewer neuropeptides than the ventral telencephalon; however, *cckb* and *cart3b* were commonly expressed in most of the dorsal telencephalic areas, except that *cckb* was absent in Dmp, whereas *cart3b* was absent in Dlv and the posterior zone of D (Dp) (Figs. 3–10, Table 2).

##### 3.1.1.3 Ventral telencephalon

The ventral telencephalic area (V) contains nine nuclei (Figs. 3–10, Table 3). In contrast to the abundant presence of glutamatergic neurons in the dorsal telencephalon, most neurons in the ventral telencephalic area were GABAergic inhibitory neurons (Figs. 3–10, Table 3). The dorsal nucleus of V (Vd) and supracommissural nucleus of V (Vs) are anteroposteriorly adjacent structures in the dorsomedial part of the ventral telencephalon and show exactly the same pattern of neuropeptide expression, implying similar functional features of these nuclei (Figs. 3–7. Table 3). The postcommissural nucleus of V (Vp) also expressed a similar repertoire of neuropeptides as Vd and Vs but additionally expressed *cart5* and *npb* (Figs. 8, 9, Table 3). The intermediate nucleus of V (Vi) is located in the most posterodorsomedial part of the ventral telencephalon, and its presence has recently been proposed in zebrafish (Biechl *et al*., 2017). Vi shared the expression of many neuropeptides with Vd, Vs, and Vp but was characterized by the lack of *crhb* and *npy* (Figs. 9 and 10, Table 3). The ventral part of the entopeduncular nucleus (ENv) and dorsal part of the entopeduncular nucleus (ENd) differed greatly in the neurotransmitter profiles: ENd was mostly occupied by GABAergic neurons (Figs. 5-8), while ENv was composed predominantly of glutamatergic neurons (Figs. 8–10). In addition, *crhb* and *npy* were selectively and strongly expressed in ENd, but not in ENv (Figs. 5–10, Table 3).

**Table 3.** The summary of gene expressions in the ventral telencephalon.

|  | Ventral telencephalon |  |  |  |  |  |  |  |  |
| --- | --- | --- | --- | --- | --- | --- | --- | --- | --- |
|  | VI | Vc | Vv | Vd | Vs | Vp | Vi | ENd | ENv |
| agrp | - | - | - | - | - | - | - | - | - |
| asip2b | - | - | - | - | - | - | - | - | - |
| avp | - | - | - | - | - | - | - | - | - |
| calca | - | - | - | - | - | - | - | - | - |
| cart2a | - | - | - | - | - | - | - | - | - |
| cart2b | - | - | - | - | - | - | - | - | - |
| cart3a | - | - | + | + | + | + | + | + | + |
| cart3b | - | - | + | + | + | + | + | + | - |
| cart4 | - | - | - | - | - | - | - | - | - |
| cart5 | - | - | + | - | - | + | - | - | + |
| ccka | - | - | - | - | - | - | - | - | - |
| cckb | - | + | - | + | + | + | + | + | - |
| crha | - | - | - | - | - | - | - | - | - |
| crhb | - | - | + | + | + | + | - | + | - |
| gal | - | - | - | - | - | - | - | - | - |
| galp | - | - | - | - | - | - | - | - | - |
| gnrh2 | - | - | - | - | - | - | - | - | - |
| gnrh3 | - | - | - | - | - | - | - | - | - |
| grp | - | - | - | - | - | - | - | - | - |
| hcrt | - | - | - | - | - | - | - | - | - |
| kiss1 | - | - | - | - | - | - | - | - | - |
| kiss2 | - | - | - | - | - | - | - | - | - |
| npb | + | - | + | - | - | + | - | - | - |
| npvf | - | - | - | - | - | - | - | - | - |
| npy | - | - | + | + | + | + | - | + | - |
| oxt | - | - | - | - | - | - | - | - | - |
| pdyn | - | - | + | + | + | + | + | - | - |
| penka | + | - | + | + | + | + | + | - | + |
| penkb | - | + | + | + | + | + | + | - | - |
| pmch | - | - | - | - | - | - | - | - | - |
| pmchl | - | - | + | - | - | - | - | - | - |
| pomca | - | - | - | - | - | - | - | - | - |
| pomcb | - | - | + | - | - | - | - | - | - |
| tac1 | - | + | + | + | + | + | + | - | - |
| tac3a | - | - | + | + | + | + | + | - | - |
| tac3b | - | - | - | - | - | - | - | - | - |
| vipa | - | - | + | - | - | - | - | - | - |
| vipb | - | - | - | - | - | - | - | - | - |
| gad mix | + | + | + | + | + | + | + | + | + |
| vglut mix | + | - | + | + | + | + | + | - | + |

#### 3.1.2 Diencephalon

##### 3.1.2.1 Preoptic area

Four nuclei were observed in the preoptic area, all of which expressed numerous neuropeptides (Figs. 7–12, Table 4). All nuclei commonly expressed *ccka*, *crhb*, *npy*, *pdyn*, *penka*, *penkb*, *tac1*, and *tac3a* (Figs. 7–12, Table 4). The anterior part of the parvocellular preoptic nucleus (PPa) expressed 21 neuropeptides among the 38 tested, with the largest number in the preoptic area (Figs. 7–9. Table 4). The dorsal and ventral parts of the PPa showed different neuropeptide expression patterns (see Section 3.2.1). The posterior part of the parvocellular preoptic nucleus (PPp) was characterized by densely packed GABAergic neurons, and its neuropeptide-expressing cells were sparser than those in other preoptic nuclei (Figs. 10–12). The magnocellular preoptic nucleus (PM) contained *avp*-, *oxt*-, *crhb*-, or *penka*-expressing magnocellular cells, as well as parvocellular cells positive for *asip2b*, *cart5*, *ccka*, *gal*, *galp*, *npy*, *pdyn*, *penkb*, *tac1*, and *tac3a* (Figs. 10 and 11). In addition, cells with very weak expressions of *crha* and *npb* were also observed (Figs. 10 and 11).

**Table 4.**
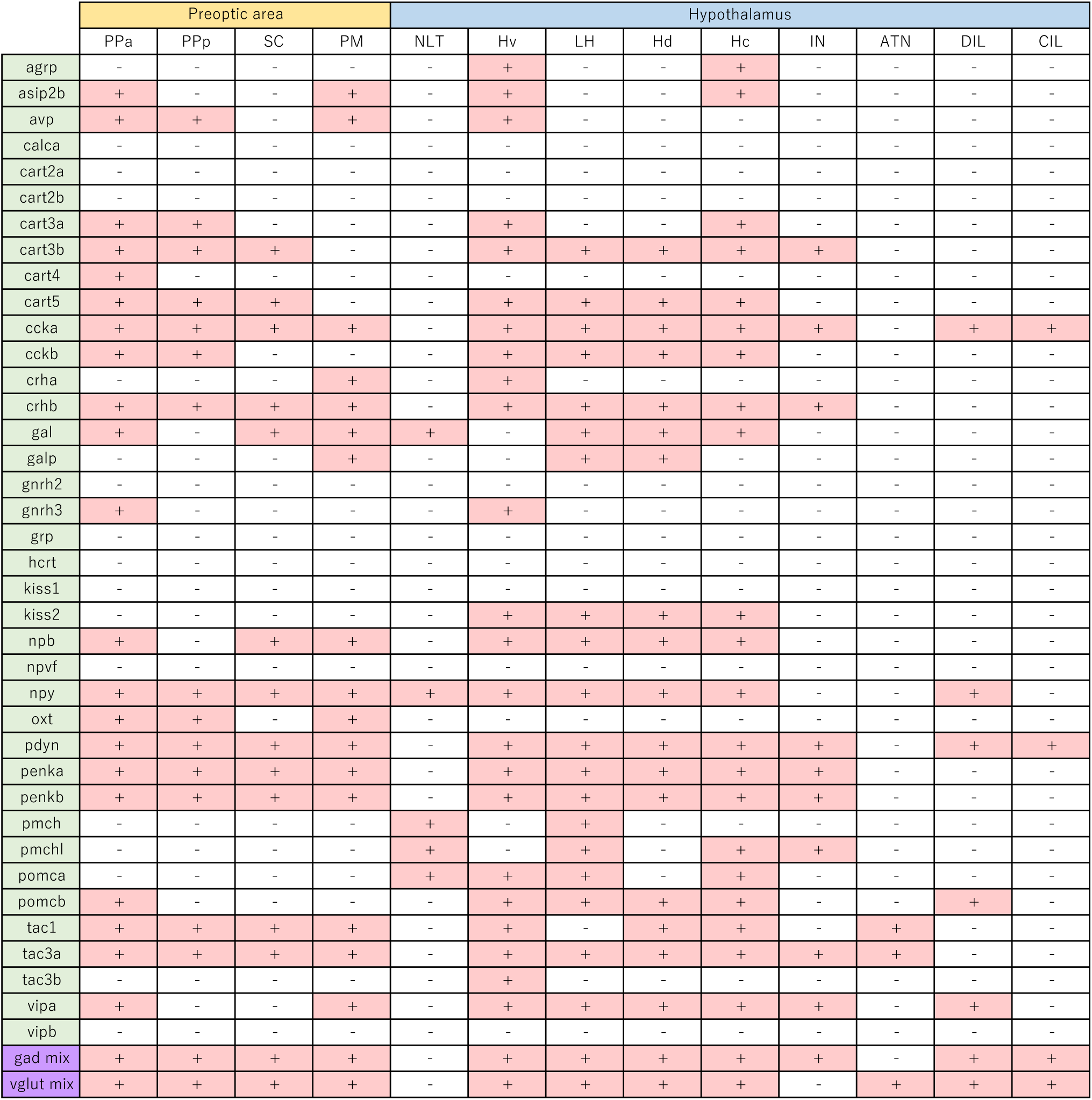
The summary of gene expressions in the preoptic area and hypothalamus.

##### 3.1.2.2 Hypothalamus

The hypothalamus contains nine nuclei (Figs. 12–20, Table 4). NLT, which is a thin cell layer at the outer rim of the hypothalamus, expressed *gal*, *npy*, *pmch*, *pmchl*, and *pomca* (Figs. 12, 13, Table 4). The ventral zone of the periventricular hypothalamus (Hv), lateral hypothalamic nucleus (LH), dorsal zone of the periventricular hypothalamus (Hd), and caudal zone of the periventricular hypothalamus (Hc) commonly expressed *cart3b*, *cart5*, *ccka*, *cckb*, *crhb*, *kiss2*, *npb*, *npy*, *pdyn*, *penka*, *penkb*, *pomcb*, *tac3a*, and *vipa* (Figs. 12–20, Table 4). Furthermore, Hv expressed 23 neuropeptides, many of which were localized ventrally (see 3.2.2) (Figs. 12–16). Neuropeptide expression in Hd and Hc was also not uniform, with distinct subregions within the nucleus that displayed different neuropeptide expression profiles (see 3.2.3 and 3.2.4) (Figs. 14–18). The IN is adjacent to the PTN and is distinguishable from it by the predominance of GABAergic neurons, while PTN neurons are glutamatergic (Figs. 15, 16). The anterior tuberal nucleus (ATN) differed significantly from other hypothalamic nuclei in that it expressed only two neuropeptides, *tac1* and *tac3a*, and contained only glutamatergic neurons (Figs. 13–16), which is a feature similar to the mammalian ventromedial hypothalamic nucleus (VMN) (see 3.3). Furthermore, the central (CIL) and diffuse (DIL) nucleus of the inferior lobe differed from other hypothalamic nuclei in the predominance of glutamatergic neurons and low neuropeptide expression (Figs. 14–20). A large part of the inferior lobe, including the CIL and DIL, has recently been shown to be of mesencephalic origin, and it has been debated whether these nuclei should be included or excluded from the hypothalamus (Bloch *et al*., 2019). Our results also suggest that these nuclei differ significantly from other hypothalamic nuclei in terms of neuropeptide expression profiles.

**Figure 17.**
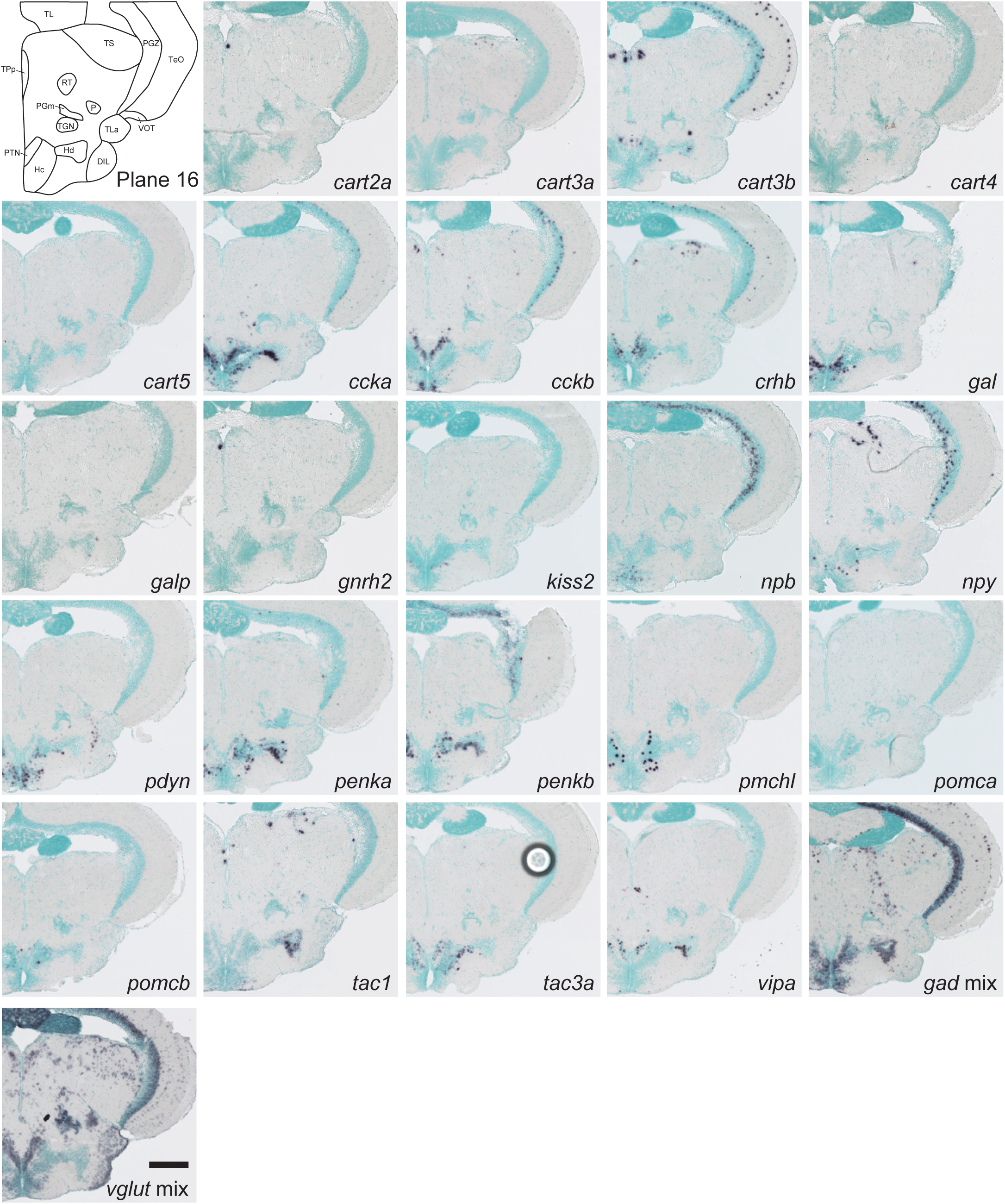
Schematic of plane 16 and photographs of ISH results of genes expressed in plane 16. Scale bar = 200 μm.

**Figure 18.**
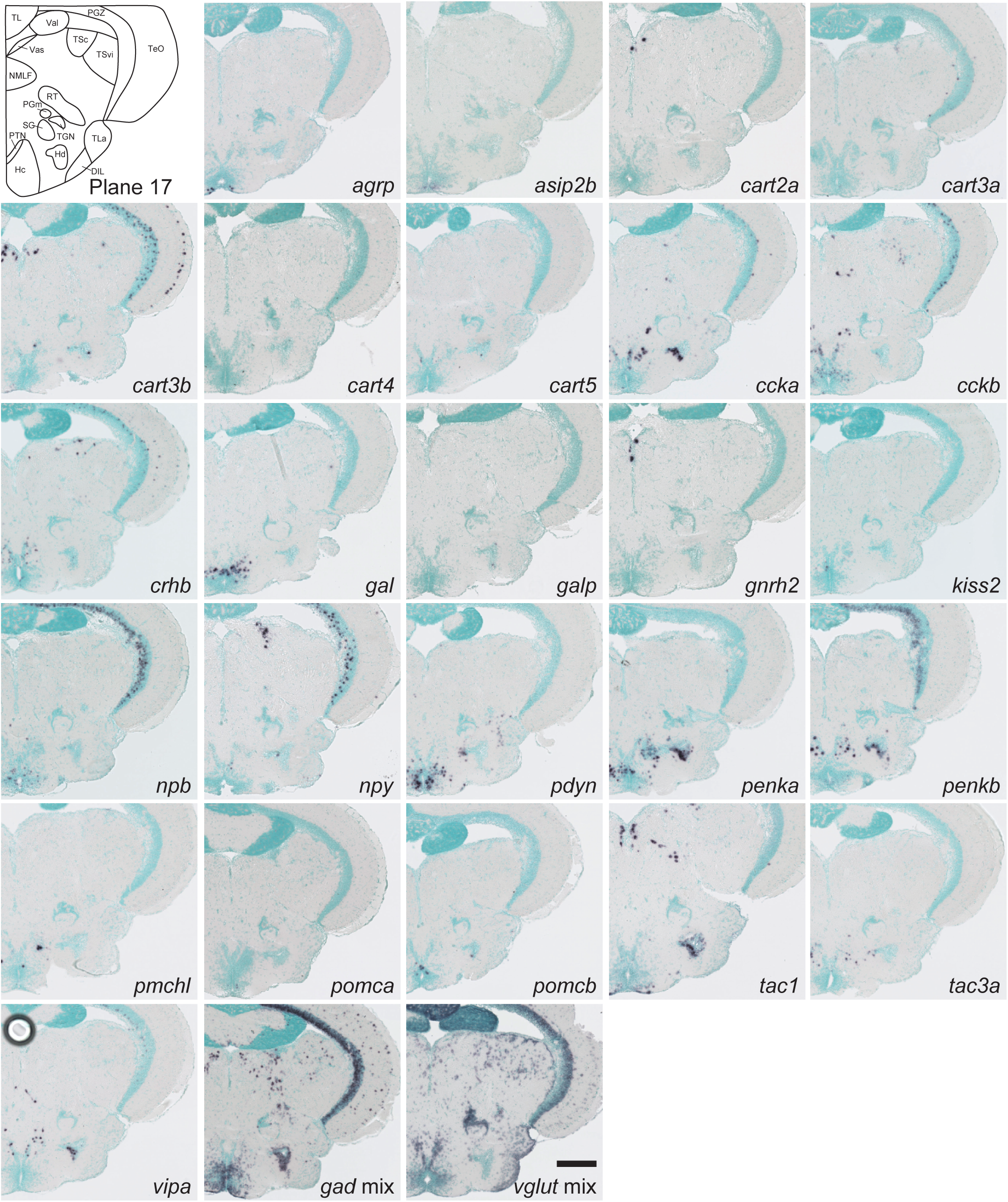
Schematic of plane 17 and photographs of ISH results of genes expressed in plane 17. Scale bar = 200 μm.

**Figure 19.**
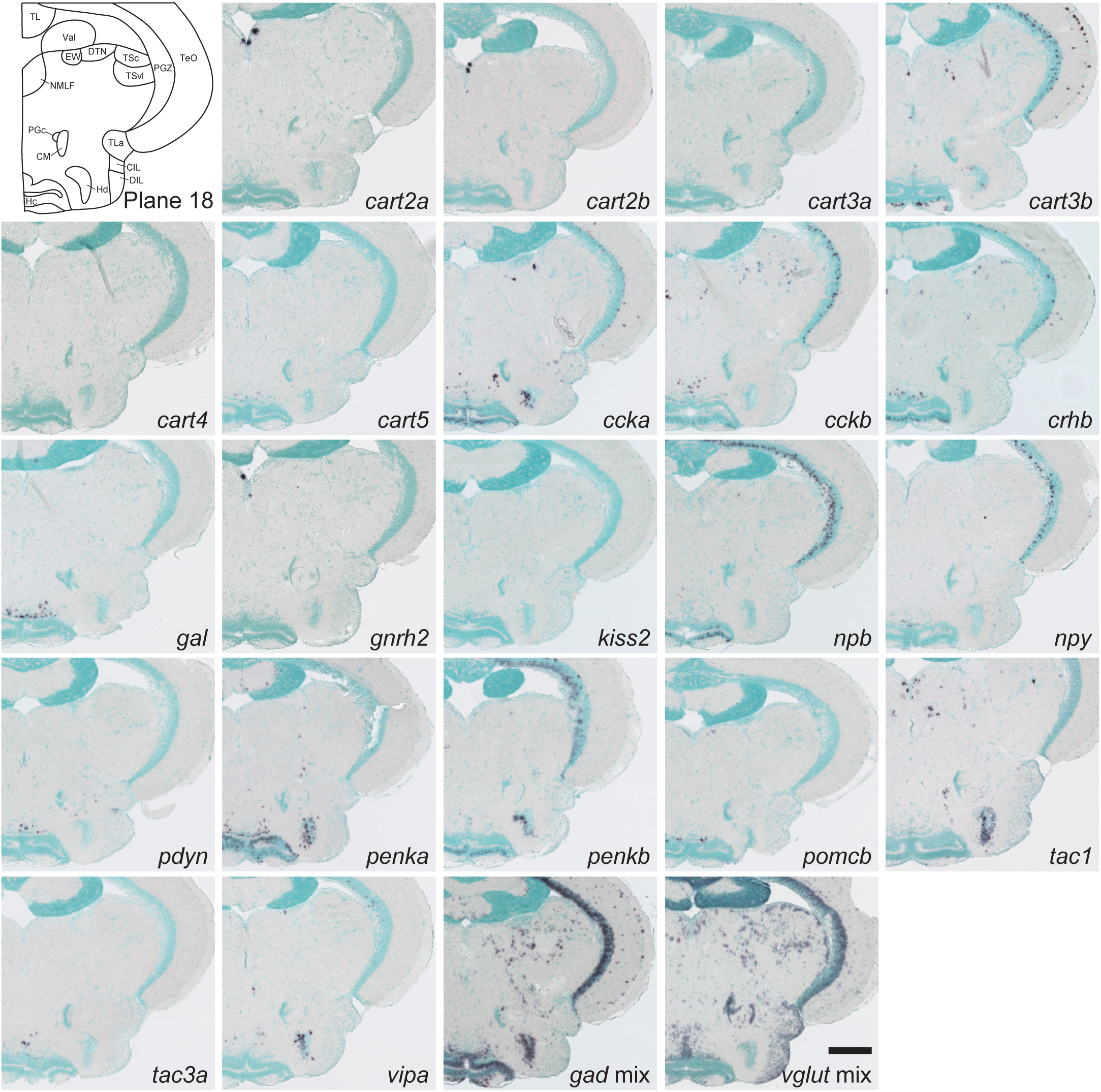
Schematic of plane 18 and photographs of ISH results of genes expressed in plane 18. Scale bar = 200 μm.

**Figure 20.**
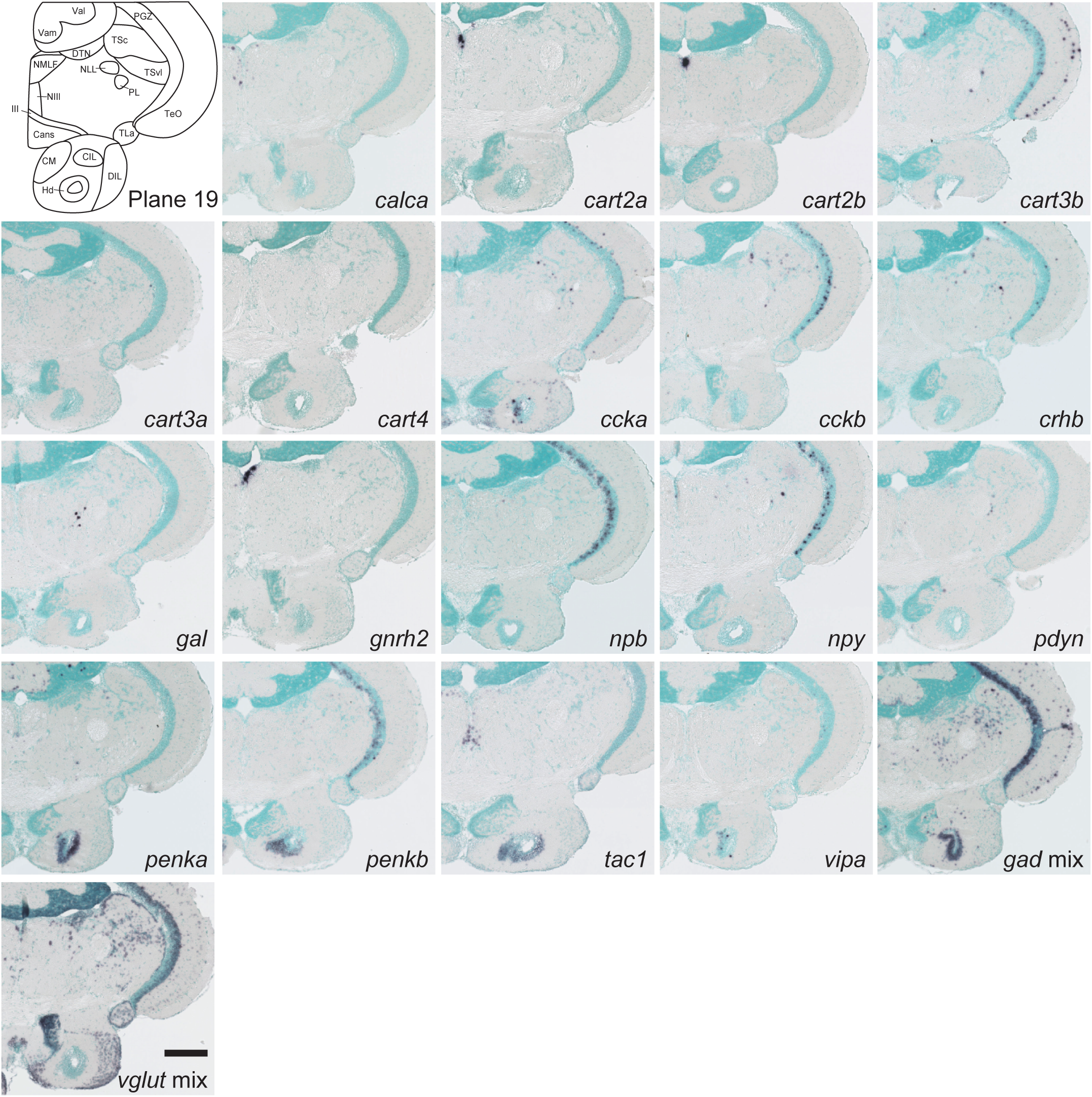
Schematic of plane 19 and photographs of ISH results of genes expressed in plane 19. Scale bar = 200 μm.

##### 3.1.2.3 Prethalamus

Prethalamus contains four nuclei (Figs. 10–13, Table 5). The rostrolateral nucleus (R) did not express any of the neuropeptides examined (Fig. 11). The intermediate thalamic nucleus (I), ventromedial thalamic nucleus (VM), and ventrolateral thalamic nucleus (VL) are located adjacent to each other; however, their neuropeptide expression patterns differed. The three nuclei commonly expressed *cart3b* and *penka* (Figs. 10–13). VM expressed seven neuropeptides, among which *npb*, *pdyn*, *vipa*, and *vipb* were present only in the VM among the prethalamic nuclei (Figs. 1–13). The VL had a higher percentage of glutamatergic neurons than the VM, and *cckb* expression was specific to the VL in the prethalamus (Figs. 11–13). *npy* and *penkb* were expressed only in I among the prethalamic nuclei (Fig. 11).

**Table 5.** The summary of gene expressions in the prethalamus, posterior tuberculum, and thalamus.

|  | Prethalamus |  |  |  | Posterior tuberculum |  |  | Thalamus |  |  |  |  |  |
| --- | --- | --- | --- | --- | --- | --- | --- | --- | --- | --- | --- | --- | --- |
|  | R | VM | VL | I | TPp | PVO | PTN | Had | Hav | A | DP | IC | CP |
| agrp | - | - | - | - | - | - | - | - | - | - | - | - | - |
| asip2b | - | - | - | - | - | - | - | - | - | - | + | - | - |
| avp | - | - | - | - | - | - | - | - | - | - | - | - | - |
| calca | - | - | - | - | + | - | - | - | - | - | - | - | - |
| cart2a | - | - | - | - | - | - | - | - | - | - | - | - | - |
| cart2b | - | - | - | - | - | - | - | - | - | - | - | - | - |
| cart3a | - | - | - | - | + | - | + | - | - | + | + | - | - |
| cart3b | - | + | + | + | + | - | + | - | - | - | + | + | - |
| cart4 | - | - | - | - | - | - | - | - | - | - | - | - | - |
| cart5 | - | - | - | - | - | - | - | - | - | - | - | - | - |
| ccka | - | - | - | - | + | + | + | - | - | - | - | + | + |
| cckb | - | - | + | - | + | + | + | + | - | + | + | - | + |
| crha | - | - | - | - | - | - | - | - | - | - | - | - | - |
| crhb | - | + | + | - | + | + | + | - | - | - | + | + | - |
| gal | - | - | - | - | - | - | - | - | - | - | - | - | - |
| galp | - | - | - | - | + | - | - | - | - | - | - | - | - |
| gnrh2 | - | - | - | - | - | - | - | - | - | - | - | - | - |
| gnrh3 | - | - | - | - | - | - | - | - | - | - | - | - | - |
| grp | - | - | - | - | - | - | - | - | - | - | - | - | - |
| hcrt | - | - | - | - | - | + | - | - | - | - | - | - | - |
| kiss1 | - | - | - | - | - | - | - | - | + | - | - | - | - |
| kiss2 | - | - | - | - | + | - | - | - | - | - | - | - | - |
| npb | - | + | - | - | + | - | + | - | - | - | - | + | - |
| npvf | - | - | - | - | - | + | - | - | - | - | - | - | - |
| npy | - | - | - | + | + | - | + | - | - | + | - | + | - |
| oxt | - | - | - | - | - | - | - | - | - | - | - | - | - |
| pdyn | - | + | - | - | - | + | + | + | - | - | - | + | - |
| penka | - | + | + | + | + | + | + | - | - | - | - | + | + |
| penkb | - | - | - | + | + | + | + | - | - | + | - | + | - |
| pmch | - | - | - | - | - | - | - | - | - | - | - | - | - |
| pmchl | - | - | - | - | - | + | - | - | - | - | - | - | - |
| pomca | - | - | - | - | - | + | - | - | - | - | - | - | - |
| pomcb | - | - | - | - | - | - | + | - | - | - | - | - | - |
| tac1 | - | - | - | - | + | - | - | + | - | - | - | - | + |
| tac3a | - | - | - | - | + | + | + | + | - | - | - | - | - |
| tac3b | - | - | - | - | + | - | - | - | - | - | - | - | + |
| vipa | - | + | - | - | + | - | + | - | - | - | - | - | - |
| vipb | - | + | - | - | - | - | - | - | - | - | - | - | - |
| gad mix | + | + | + | + | + | + | + | - | - | + | - | + | - |
| vglut mix | + | + | + | + | + | + | + | + | + | + | + | + | + |

##### 3.1.2.4 Posterior tuberculum

The periventricular nucleus of the posterior tuberculum (TPp), PVO, and PTN in the posterior tuberculum, which are known to contain many dopaminergic neurons (Rink and Wullimann, 2001), also showed expression of multiple neuropeptides (Figs. 13–18, Table 5). Of note is the specific expression of *calca* only in the TPp of the forebrain (Figs. 14 and 15). Furthermore, *hcrt* and *npvf* were selectively localized to the PVO in the forebrain (Figs. 14 and 15).

##### 3.1.2.5 Thalamus

The zebrafish thalamus has seven nuclei (Figs. 10–14, Table 5). The habenula is a conserved structure among all vertebrates. In teleost, it is divided into the dorsal (Had) and ventral (Hav) habenular nuclei, which differ in neuronal connectivity and function (Okamoto *et al*., 2012). Neuropeptide expression also differed between Had and Hav, with *kiss1* being the only neuropeptide expressed in Hav (Figs. 11–13). Had can be further divided into Hadm and Hadl (Agetsuma *et al*., 2010). *pdyn* and *tac1* were expressed in Hadl, whereas *tac3a* was expressed in Hadm (Figs. 11 and 12). The dorsal posterior thalamic nucleus (DP) was the only nucleus other than the hypothalamus expressing *asip2b* (Fig. 16). IC was recently defined based on its high expression in *gad* (Mueller and Guo, 2009). Strong expression of *npy* and *penkb* also distinguished this nucleus from other adjacent nuclei (Figs. 14,15).

##### 3.1.2.6 Preglomerular complex

Neuropeptide expression was low in the nuclei of the preglomerular complex. The subglomerular nucleus (SG), tertiary gustatory nucleus (TGN), anterior preglomerular nucleus (PGa), caudal preglomerular nucleus (PGc), and corpus mamillare (CM) did not express any of the neuropeptides examined (Figs. 13–20, Table 6). In contrast, a few neuropeptides were expressed in the lateral (PGl) and medial (PGm) preglomerular nuclei. In addition, the torus lateralis (TLa) and posterior thalamic nucleus (P) only expressed *tac1* (Figs. 15–17).

**Table 6.**
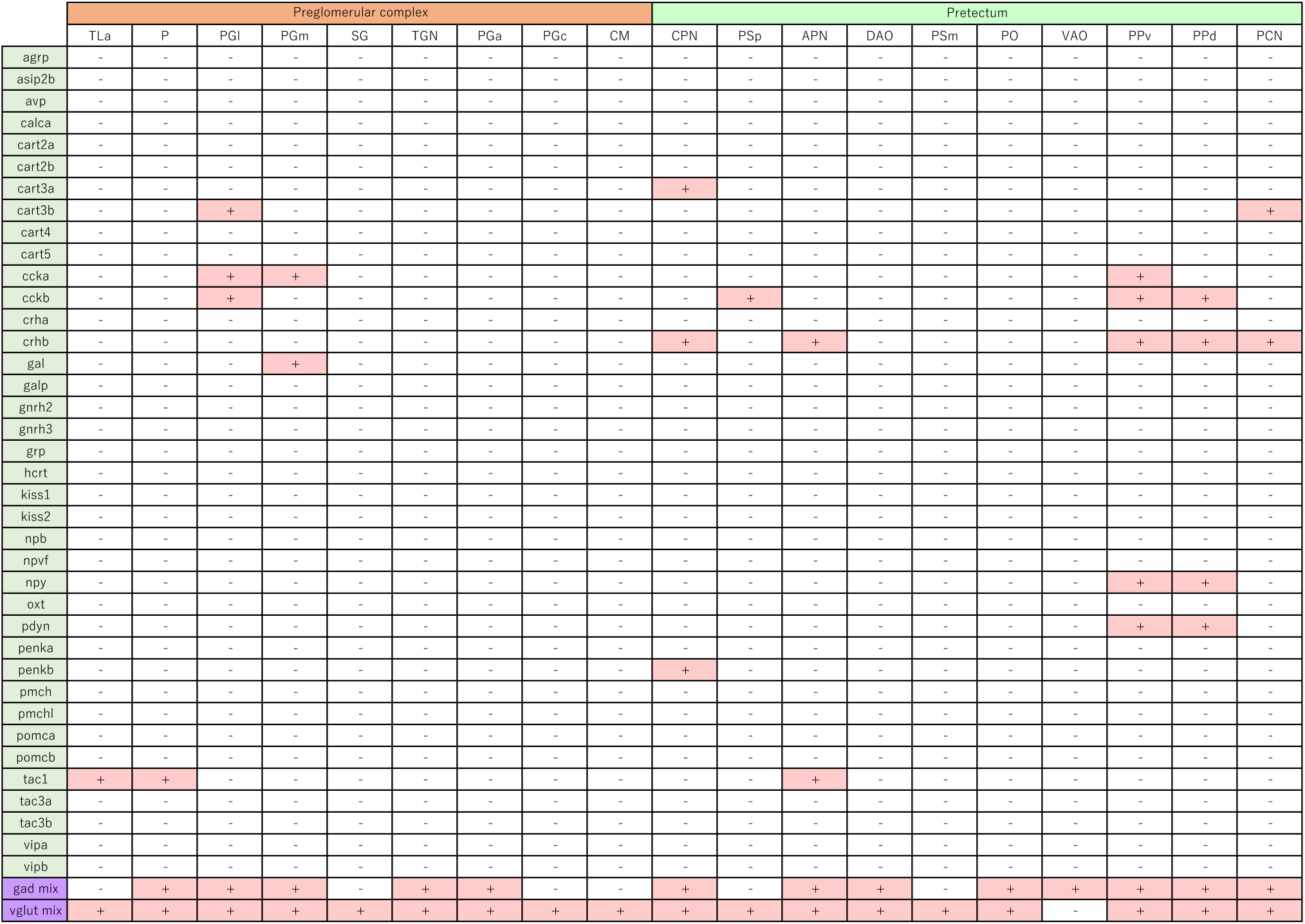
The summary of gene expressions in the preglomerular complex and pretectum.

##### 3.1.2.7 Pretectum

Neuropeptide expression was low in the nuclei of the pretectum (Figs. 10–14, Table 6). The dorsal accessory optic nucleus (DAO), magnocellular superficial pretectal nucleus (PSm), posterior pretectal nucleus (PO), and ventral accessory optic nucleus (VAO) did not express the examined neuropeptides. The remaining nuclei expressed neuropeptides with distinct expression patterns.

#### 3.1.3 Individual differences

For each gene, we examined the brains of 3–5 male zebrafish and found that, in most cases, the expression patterns were consistent across individuals (although not compared quantitatively). Therefore, only representative pictures from one fish for each gene are shown in Figs. 2–20. In contrast, several genes showed marked individual differences in their expression patterns, as summarized in Supplemental Fig. 2. *asip2b* in PPa was observed in only two of the four fish examined. *asip2b* in DP was also observed in only two of the four fish examined. As *asip2b* (also referred to as *agrp2*) has been reported to play a role in background adaptation (Zhang *et al*., 2010) and stress axis regulation (Shainer *et al*., 2019), these individual differences might be related to these factors. *agrp* was expressed in Hv in all four fish examined; however, the signal densities were highly variable. As for *pomca*, in four fish examined, one showed expression in both NLT and Hv, while the other three showed expression only in Hv. As *agrp* and *pomca* expression is known to be regulated by feeding conditions (*agrp*, Jeong *et al*., 2018; *pomca*, Löhr *et al*., 2018), these individual differences may reflect different feeding statuses.

### 3.2 Novel subnuclei revealed by neuropeptide expressions in the hypothalamus

Neuropeptides are prominently expressed in the hypothalamus and are, therefore, considered to play crucial roles in various hypothalamic functions. Our scrutinized observation of the expression patterns of neuropeptides in the hypothalamus revealed that within some of the hypothalamic nuclei defined by the zebrafish brain atlas (Wullimann *et al*., 1996), there were distinct subnuclei that were not annotated, but displayed distinct neuropeptide expression patterns.

#### 3.2.1 Subnclei in the PPa

The anterior part of the parvocellular preoptic nucleus (PPa) is a relatively large region covering all of the anterior preoptic area (Fig. 21 A). Within the PPa, there is a dense mass of neurons in the dorsal part of the PPa (PPad), segregated from the ventral part of the PPa (PPav) by a thin blank area with few neurons (Fig. 21, see the counterstain). The expressions of four neuropeptides (*cart5*, *ccka*, *gnrh3*, and *vipa*) were restricted to the PPav (Figs. 7-9), as represented by *ccka* and *vipa* in Fig. 21. Although other neuropeptides present in PPa were expressed in both PPad and PPav, their expression patterns were not uniform: some genes such as *gal* and *pdyn* were expressed predominantly in PPav, while others such as *cart3b* and *tac3a* were expressed more prominently in PPad (Fig. 21). In addition, glutamatergic neurons were mostly restricted to PPav, whereas GABAergic neurons were present in both PPad and PPav (Fig. 21). Thus, based on the expression patterns of neuropeptides/neurotransmitters, we proposed two subnuclei in PPa: PPad and PPav.

**Figure 21.**
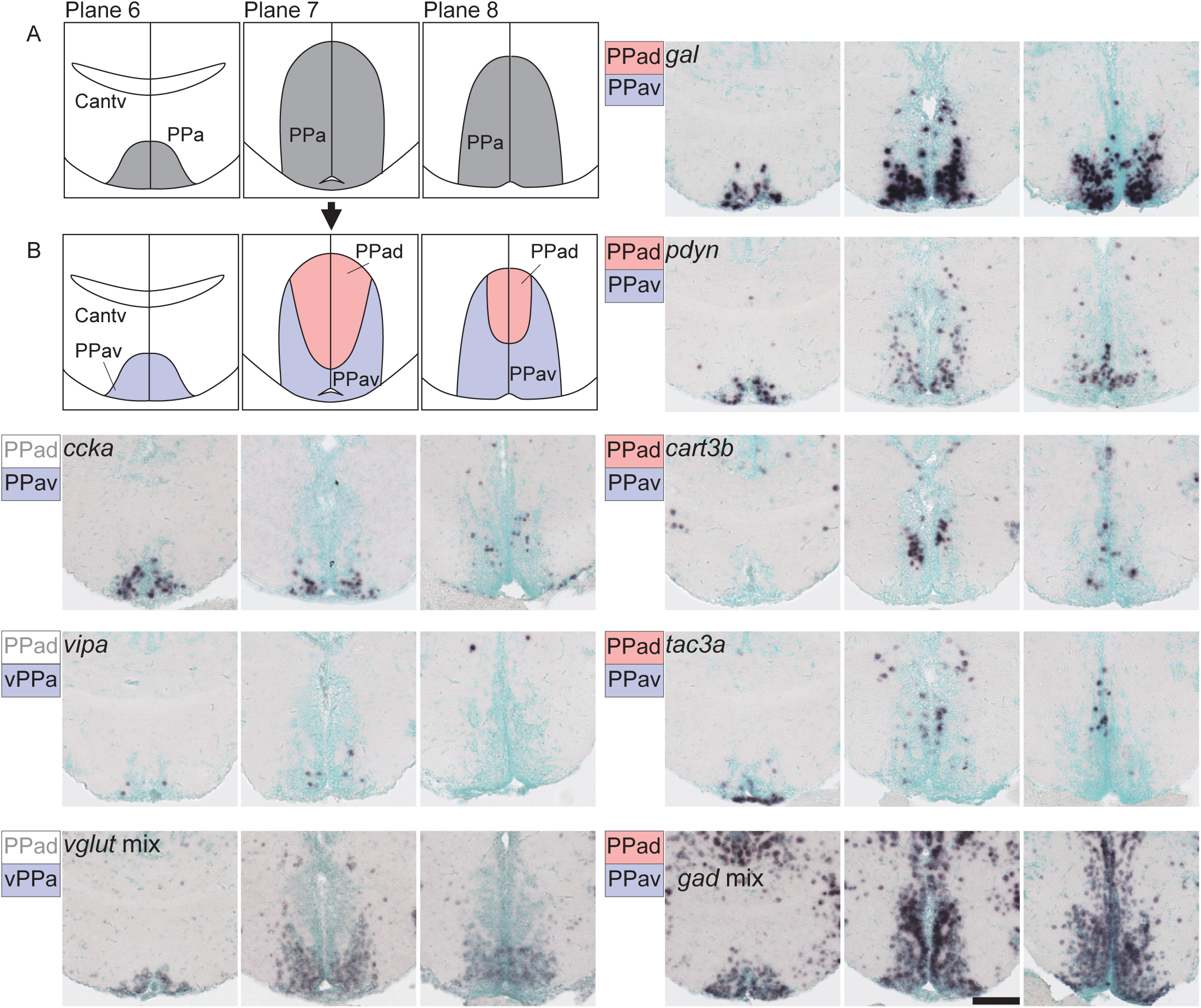
Subnuclei in the anterior part of the parvocellular preoptic nucleus (PPa). (A) The PPa (grey) in the original zebrafish brain atlas. (B) The PPa can be divided into two subnuclei: the ventral part of PPa (PPav, pink) and dorsal part of PPa (PPad, purple). Neuropeptides such as *ccka* and *vipa* and *vglut* mix are exclusively expressed in the PPav. *gal* and *pdyn* are expressed in both the PPad and PPav, but predominantly in the PPav. *cart3b* and *tac3a* are also expressed in the PPav and PPad, but predominantly in the PPad. *gad* mix is expressed in both the PPad and PPav. Scale bar = 100 μm.

#### 3.2.2 Subnuclei in Hv

The ventral zone of the periventricular hypothalamus (Hv) is a layer of neurons surrounding the diencephalic ventricle (DiV) (Fig. 22). Most of the neuropeptides present in the Hv (*agrp*, *asip2b*, *cart3b*, *cart3a*, *ccka*, *crha*, *crhb*, *gnrh3*, *npb*, *penka*, *penkb*, *pomcb*, and *vipa*) were expressed exclusively in the ventral part of Hv (Hvv) (Figs. 12-16), as represented by *agrp* and *ccka* in Fig. 22. In contrast, four genes (*pdyn*, *tac1*, *tac3a*, and *tac3b*) were expressed in the dorsal part of Hv (Hvd) (Figs. 12-16) as represented by *pdyn* and *tac3a* in Fig. 22. The boundary between Hvd and Hvv was clearly defined by *pdyn* expression (Fig. 22 B). The genes expressed in Hvd and Hvv were mutually exclusive, with only one exception: *cckb*, which was highly expressed in Hvv and slightly expressed in the anterior part of Hvd (Fig. 22). Hvd and Hvv also differed in neurotransmitter expression patterns: Hvd contained mostly glutamatergic neurons, whereas Hvv harbored mostly GABAergic neurons (Fig. 22). Thus, we propose that Hv can be divided into two subnuclei, Hvd and Hvv.

**Figure 22.**
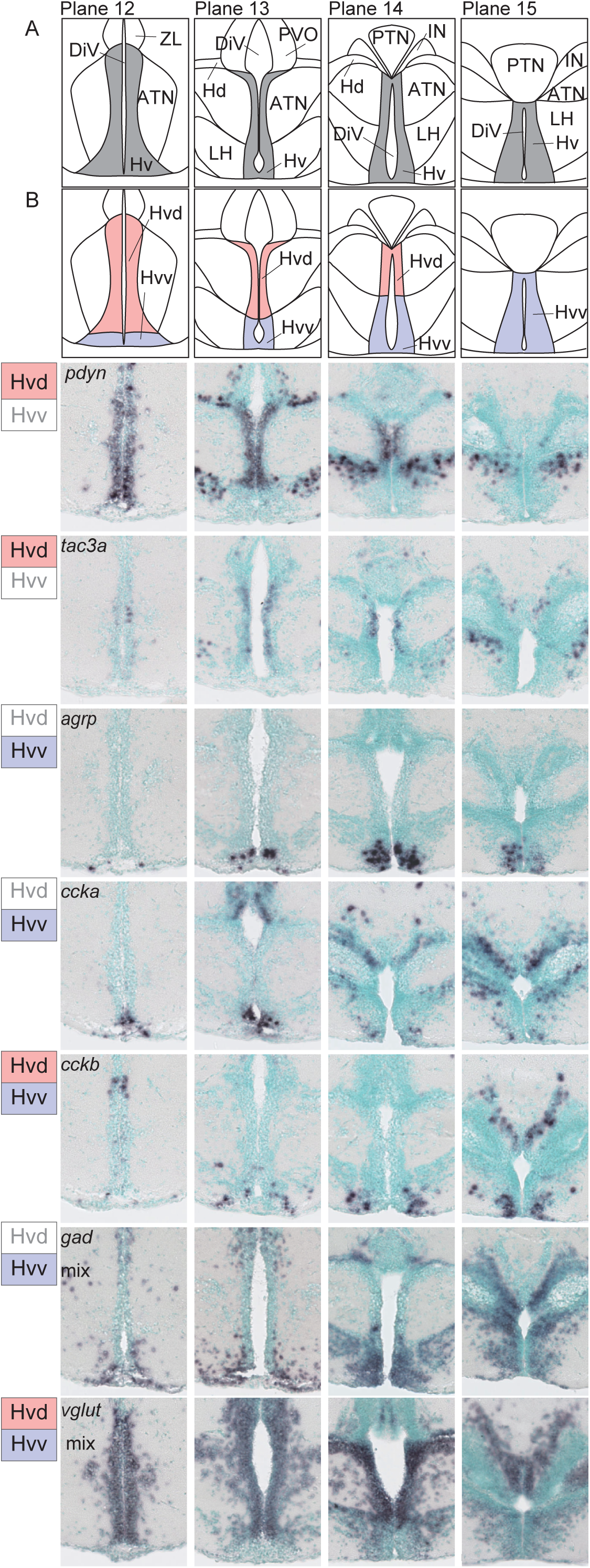
Subnuclei in the ventral zone of the periventricular hypothalamus (Hv). (A) The Hv (grey) in the original zebrafish brain atlas. (B) The Hv can be divided into two subnuclei: the ventral part of Hv (Hvv, purple) and the dorsal part of Hv (Hvd, pink). Neuropeptides such as *pdyn* and *tac3a* are expressed only in the Hvd, whereas *agrp* and *ccka* are expressed only in the Hvv. *cckb* is expressed in both the Hvd and Hvv. *gad* mix is predominantly expressed in the Hvv and the *vglut* mix is more densely expressed in the Hvd. Scale bar = 100 μm.

#### 3.2.3 Subnuclei in Hd

The dorsal zone of the periventricular hypothalamus (Hd) is the nucleus that surrounds the lateral recess of the DiV (LR). At the anterior end (Fig. 14; plane 13 in Fig. 23), Hd was located medially, adjacent to Hv. More posteriorly (Fig. 15; plane 14 in Fig. 23), Hd extends laterally and continues caudally through the lateral part of the hypothalamus. The expression of neuropeptide genes differed along the mediolateral and dorsoventral axes of Hd (Fig. 23). Genes such as *cart5*, *gal*, *kiss2*, *npy*, *pdyn,* and *tac3a* were expressed only in the medial part of Hd (Hdm) (Figs. 14 and 15), as represented by *npy* in Fig. 23. Within the lateral part of the Hd, the neuropeptide expression patterns were different between the dorsal and ventral parts. *cart3b* and *npb* were only expressed in the dorsolateral part of Hd (Hddl) (Figs. 15–20), as represented by *cart3b* in Fig. 23. In contrast, *ccka*, *cckb,* and *pomcb* were only expressed in the ventrolateral part of Hd (Hdvl) (Figs. 15–20), as represented by *ccka* in Fig. 23. The boundary between Hddl and Hdvl is clearly delineated by *ccka* expression. Five genes (*galp*, *penka*, *penkb*, *tac1*, and *vipa*) were expressed in all the Hd subnuclei (Figs. 14–20), as shown by *penka* and *vipa* in Fig. 23. Both Hddl and Hdvl were *gad*-positive but not *vglut*-positive (Fig. 23), whereas Hdm harbored both *gad*-positive and *vglut*-positive neurons. Altogether, we propose that Hd be divided into three subnuclei: Hdm, Hddl, and Hdvl.

**Figure 23.**
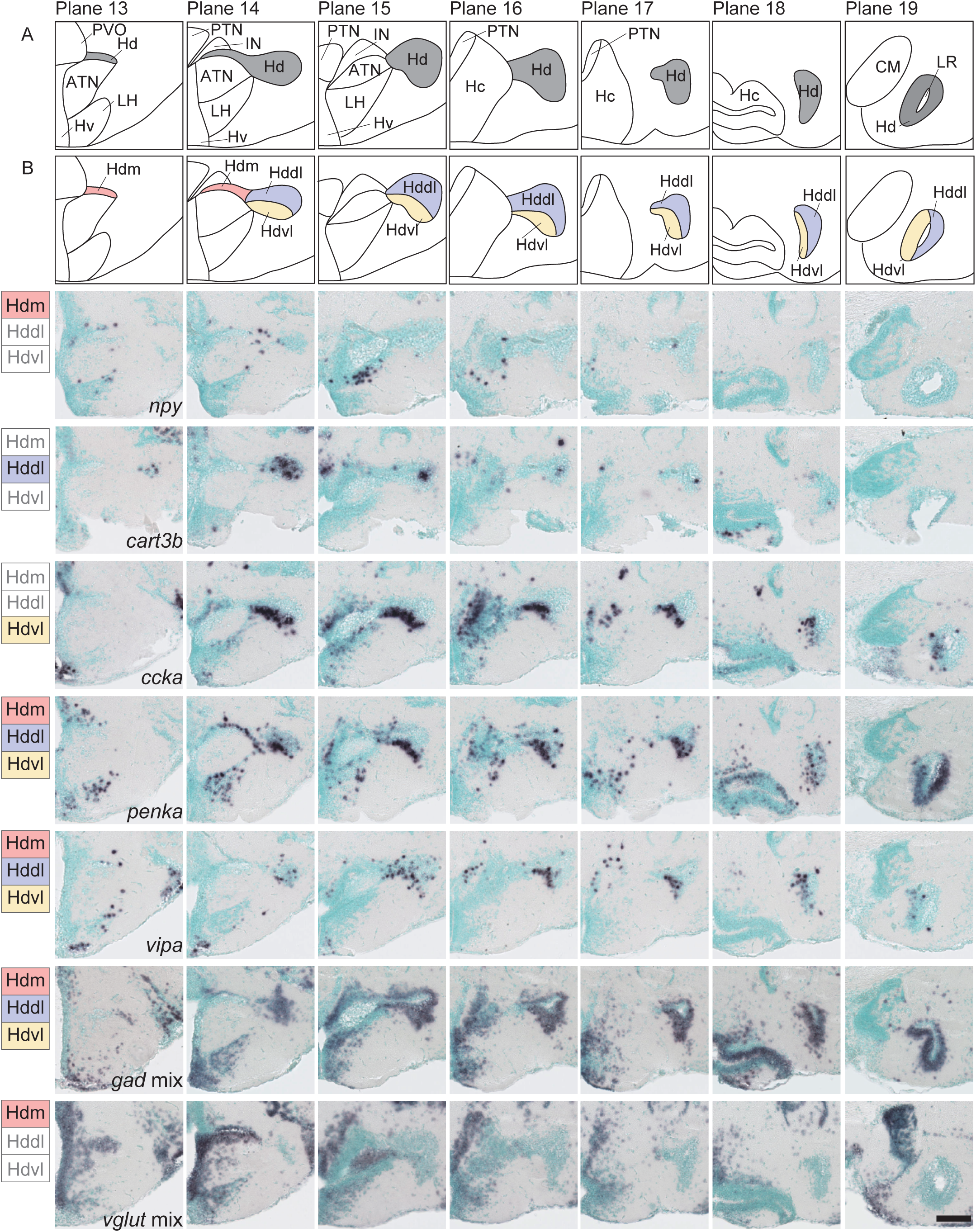
Subnuclei in the dorsal zone of the periventricular hypothalamus (Hd). (A) The Hd (grey) in the original zebrafish brain atlas. (B) The Hd can be divided into three subnuclei: the medial (Hdm, pink), dorsolateral (Hddl, purple), and ventrolateral (Hdvl, yellow) part of Hd. Some neuropeptides are expressed solely in the Hdm, Hddl, or Hdvl (represented by *npy*, *cart*, and *ccka*, respectively). Neuropeptides such as *penka* and *vipa* are expressed in all three subnuclei. *gad* mix is also expressed in all subnuclei. *vglut* mix is absent in all subnuclei. Scale bar = 100 μm.

#### 3.2.4 Subnuclei in Hc

The caudal zone of the periventricular hypothalamus (Hc) refers to the hypothalamic area caudal to the Hv and LH (Fig. 24). The anterior part of Hc is not recessed, while the posterior part surrounds the posterior recess (PR) (Fig. 24). The expression of neuropeptides in the anterior and posterior parts of Hc was distinct. In the anterior part of Hc, seven genes (*cart5*, *kiss2*, *npy*, *penkb*, *pmchl*, *pomcb*, and *tac3a*) were expressed only in the dorsal part (the anterodorsal part of Hc, Hcad) (Figs. 17, 18), as shown by *pomcb* and *tac3a* in Fig. 24. Underneath Hcad, there was a triangular dense cell population (the anterointermediate part of Hc, Hcai). *npb* and *cckb* were expressed only in Hcai (Figs. 17 and 18), as represented by *npb* in Fig. 24. *agrp* and *tac1* were expressed in the ventral cell population (the anteroventral part of Hc, Hcav).

**Figure 24.**
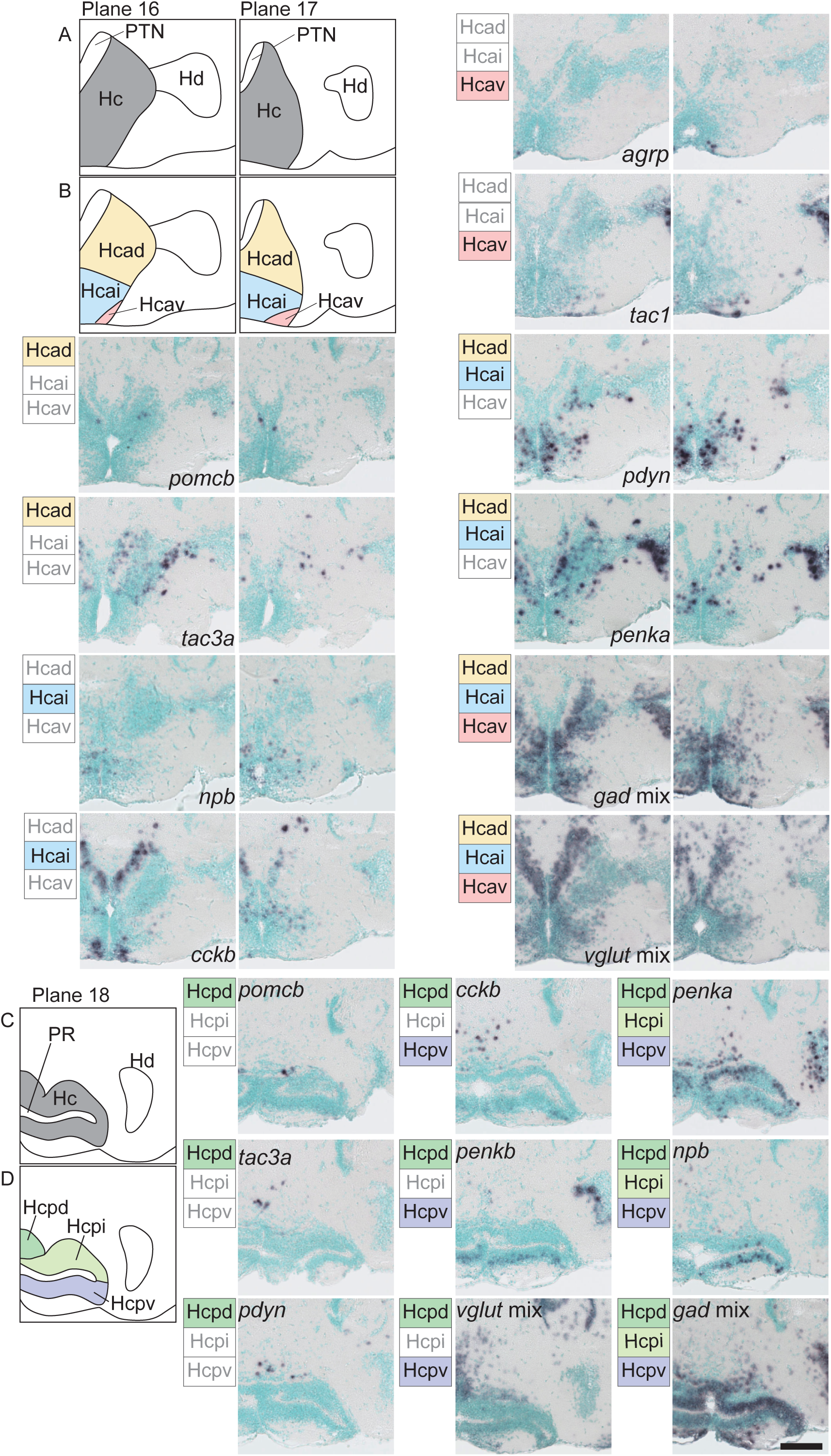
Subnuclei in the caudal zone of the periventricular hypothalamus (Hc). (A) The anterior part of Hc (grey) in the original zebrafish brain atlas. (B) The anterior part of Hc can be divided into three subnuclei: the anterodorsal (Hcad, yellow), anterointermediate (Hcai, blue), anteroventral (Hcav, pink) part of Hc. Neuropeptides such as *pomcb* and *tac3a* are solely expressed in the Hcad. Neuropeptides such as *npb* and *cckb* are expressed exclusively in the Hcai. *agrp* and *tac1* are only expressed in the Hcav. Neuropeptides such as *cckb*, *pdyn*, and *penka* are expressed across multiple subnuclei. *gad* mix and *vglut* mix are expressed in all three subnuclei. (C) Posterior part of Hc (grey) in original zebrafish brain atlas. (D) The posterior part of Hc can be divided into three subnuclei: the posterodorsal (Hcpd, green), posterointermediate (Hcpi, light green), posteroventral (Hcpv, purple) part of Hc. Neuropeptides such as *pomcb*, *tac3a*, and *pdyn* are solely expressed in the Hcpd. *cckb*, *penkb*, and *vglut* mix are expressed in the Hcpd and Hcpv. *penka*, *npb*, and *gad* mix are expressed in all three subnuclei. Scale bar = 100 μm.

The posterior part of the Hc was divided into three regions. The dense neuronal population surrounding the PR was divided into two parts: the subregion dorsal to the PR (posterointermediate part of the Hc, Hcpi) and the subregion ventral to the PR (posteroventral part of the Hc, Hcpv). Dorsal to the Hcpi is another subregion with a lower cell density (the posterodorsal part of Hc, Hcpd). These three subregions showed different neuropeptide expression patterns. Nine genes were expressed only in Hcpd (*cart5*, *gal*, *kiss2*, *npy*, *pdyn*, *pomcb*, *tac1*, *tac3a*, and *vipa*) (Fig. 19), as represented by *pomcb*, *tac3a,* and *pdyn* in Fig. 24. Three genes were expressed in Hcpd and Hcpv (*cart3b*, *cckb*, and *penkb*) (Fig. 19), as represented by *cckb* and *penkb* in Fig. 24. Four genes (*ccka*, *crhb*, *npb*, and *penka*) were expressed in all three regions (Fig. 19), as shown by *penka* and *npb* in Fig. 24. Genes expressed in both Hcpi and Hcpv were not uniformly expressed; all neuropeptides, except *penka*, were more abundantly expressed in Hcpv. *vglut* was expressed in Hcpd and in some cells in Hcpv, but not in Hcpi. *gad* was expressed in all three subregions. Thus, based on the differential expression patterns of neuropeptide genes, we propose the delineation of six subnuclei in the Hc: Hcad, Hcai, Hcav, Hcpd, Hcpi, and Hcpv.

### 3.3 Comparison of neuropeptide expressions in the hypothalamus between zebrafish and mouse

Using the data obtained in this study, we next compared the neuropeptide expression profiles in the hypothalamic nuclei of zebrafish and mice (Fig. 25). Hypothalamic nuclei were hierarchically clustered based on the expression matrix. Several hypothalamic nuclei in zebrafish were highly clustered with those in mice and showed high similarity in neuropeptide expression. These include the zebrafish Hvd/ATN with mouse ventromedial hypothalamic nucleus (VMN), zebrafish PPav/PM with mouse paraventricular hypothalamic nucleus (Pa), and zebrafish PPp with mouse medial preoptic area (MPA) (Fig. 25). Both zebrafish Hvd and ATN had only one gene that differed from the mouse VMN in either the presence (*Cck*) or absence (*Pdyn*) of expression, respectively. The expression profiles of zebrafish PM and mouse Pa were mostly identical, except that *Galp* and *Vip* were expressed only in zebrafish PM. The zebrafish PPav also showed similar expression patterns to the mouse Pa, but *Vip* and *Pomc* were only expressed in the PPav. In addition, zebrafish PPav contained high numbers of both GABAergic and glutamatergic neurons, whereas mouse Pa predominantly contained glutamatergic neurons. The zebrafish PPp and mouse MPA were also remarkably similar, except that *Vip* and *Gal* were expressed only in mouse MPA.

**Figure 25.**
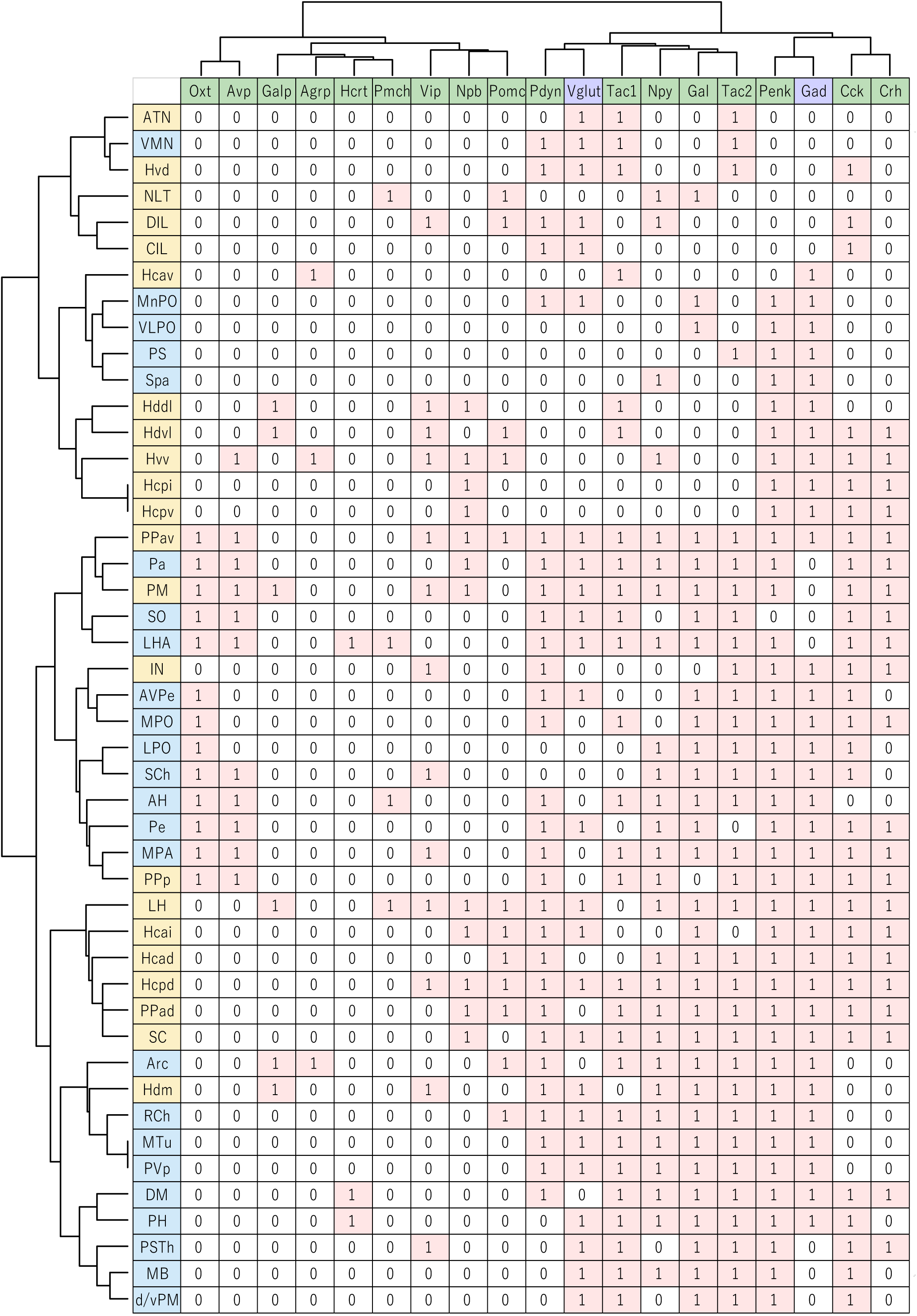
Comparison between zebrafish and mouse hypothalamic nuclei based on their neuropeptide expression patterns. Expression of neuropeptides (green) and neurotransmitter markers (purple) in the hypothalamic nuclei of zebrafish (light yellow) and mice (light blue) were extracted from the present ISH results (zebrafish), the Allen Mouse Brain Atlas, and previous papers (mouse) and summarized in a table (expressed = 1, not expressed = 0). Ward’s method hierarchically clustered zebrafish and mouse nuclei and expressed genes.

## 4. Discussion

### 4.1 Mapping of neuropeptide and neurotransmitter marker gene expressions

The expression of 38 neuropeptide genes and GABAergic/glutamatergic neuron marker genes was mapped throughout the adult zebrafish forebrain. This study presents the first large-scale atlas and database of gene expression in the adult zebrafish brain. Although expression of some of the neuropeptide genes was previously analyzed by *in situ* hybridization in the adult zebrafish brain in separate papers (*agrp*, Jeong *et al*., 2018; *avp*, Carreño Gutiérrez *et al*., 2019; *cart2b*, Akash *et al*., 2014; *cart3a*, Akash *et al*., 2014; *cart3b*, Akash *et al*., 2014; *cart5* Akash *et al*., 2014; *crha*, Grone, and Maruska, 2015; *crhb*, Alderman and Bernier, 2007; *gnrh2*, Steven *et al*., 2003; *gnrh3*, Steven *et al*., 2003; *kiss1*, Servili *et al*., 2011; *kiss2*, Servili *et al*., 2011; *hcrt*, Appelbaum *et al*., 2009; *npy*, Jeong *et al*., 2018; *npvf*, Marvel *et al*., 2018; *oxt*, Carreño Gutiérrez *et al*., 2019; *pmch*, Berman *et al*., 2009; *pmchl*, Berman *et al*., 2009; *tac1*, Ogawa *et al*., 2012; *tac3a*, Biran *et al*., 2012; Ogawa *et al*., 2012; *tac3b*, Ogawa *et al*., 2012; *gad1b*, Mueller *et al*., 2009), it has been difficult to perform detailed comparisons due to different sampling and staining conditions. The gene expression atlas is a fundamental resource in neuroscience research and will be useful for a variety of studies, including searching for marker genes for groups of neurons of interest, detailed anatomical analysis of each brain area, and investigation of homologous relationships with other species. The latter two points were further analyzed in the present study.

### 4.2 Subnuclei in the hypothalamus

The hypothalamus plays an important role as a central hub that controls the autonomic nervous system, endocrine system, and behavior. However, in teleost species, the anatomical relevance of the hypothalamic nuclei to these functions remains unclear. In addition, the definition of nuclei in the zebrafish hypothalamus is relatively simple compared with that in mammals. The zebrafish hypothalamus is divided into ten nuclei in the commonly used brain atlas (Wullimann *et al*., 1996), whereas the mouse hypothalamus is divided into over 25 nuclei in the brain atlas (Paxinos and Franklin, 2019). In this study, we attempted to define “subnuclei” in the zebrafish hypothalamus based on differential neuropeptide and neurotransmitter expression profiles to further increase the resolution of the current atlas. The information on these subnuclei defined in the present study will provide useful insights for future research on the neuroanatomy and functions of the teleost hypothalamus.

#### 4.2.1 The anterior part of the parvocellular preoptic nucleus (PPa)

The neuropeptide expression patterns revealed that PPa could be clearly divided into two subnuclei: PPad and PPav. Previous studies have shown that the expression of genes other than neuropeptides is also different between PPad and PPav, although differences in preparation make it difficult to compare the exact location of expression. Estrogen receptor 1 (*esr1*) and androgen receptor (*ar*) are mostly expressed in PPav (*esr1*, Diotel *et al*., 2011; *ar*, Gorelick *et al*., 2008). Cholinergic neurons are predominantly localized to PPad (Clemente *et al*., 2004), whereas dopaminergic neurons are confined to PPav (Yamamoto *et al*., 2011). Orthopedia a (*otpa*), a transcription factor important for the development of neurosecretory neurons, is also found only in PPav (Eugenin von Bernhardi *et al*., 2022). Recently, PPa has been shown to be critical in zebrafish social behavior (Nunes *et al*., 2021; Tallafus *et al*., 2022). PPad and PPav may play different roles in social information processing.

#### 4.2.2 The ventral zone of the periventricular hypothalamus (Hv)

Based on the expression patterns of neurotransmitters and neuropeptides, we proposed two subnuclei in Hv: Hvd and Hvv. The Hv in the teleost (also referred to as the NLTv, ventral part of the lateral tuberal nucleus) is considered homologous to the mammalian arcuate nucleus (Arc), based on the shared expression of feeding-related neuropeptide genes such as *agrp*, *pomc*, *npy*, and *cart* (Forlano and Cone, 2007; Soengas *et al*., 2018). All of these feeding-related neuropeptides are expressed exclusively in the Hvv, but not in the Hvd, suggesting a role for Hvv in the control of feeding. However, as Hvv also expresses many other neuropeptide genes, such as *ccka*/*b*, *crha*/*b*, *npb*, *penka*/*b*, and *vipa*, its function may not be confined to feeding. The function of Hvd is not yet clear, but its neuropeptide and neurotransmitter expression patterns resemble those of the mammalian ventromedial hypothalamus (VMN), as discussed below.

#### 4.2.3 The dorsal zone of the periventricular hypothalamus (Hd)

Hd can be divided into three subnuclei, Hdm, Hddl, and Hdvl. Recently, it has been reported that Hd is activated by feeding behavior (Wee *et al*., 2018) or food-related cues (Muto *et al*., 2017; Wakisaka *et al*., 2017). However, the subnuclei in the Hd that are activated and the types of neurons that are involved in feeding remain unknown. The neuropeptide expression atlas of the present study will help to clarify these important questions in the near future.

#### 4.2.4 The caudal zone of the periventricular hypothalamus (Hc)

Our analysis revealed that Hc could be divided into six subnuclei: Hcad, Hcai, Hcav, Hcpd, Hcpi, and Hcpv. The posterior region of the Hc has been reported to harbor cerebrospinal fluid (CSF)-contacting cells that produce dopamine and/or serotonin, which are absent in mammals (Xavier *et al*., 2017). Within the posterior part of the Hc, the expression patterns of neuropeptides varied greatly among the three subnuclei, suggesting that CSF-contacting cells in the posterior Hc are not uniform. In addition, the colocalization of two dopamine-synthesizing enzymes, tyrosine hydroxylase (TH) 1 and TH 2, is confined to Hcpv (Yamamoto *et al*., 2011). Functional differences in the Hc subnuclei have also been suggested in a previous study with larval zebrafish, in which the Hcpi and Hcpv responded differently to the visual cues of conspecifics (Tunbak *et al*., 2020). These regions might play distinct roles in the social reward system.

### 4.3 Comparison of hypothalamic neuropeptide expression patterns between zebrafish and mouse

To date, few studies have compared the structure of the teleost hypothalamus with that of other vertebrate lineages, and homologous relationships are not yet clear. In the present study, a comprehensive and detailed examination of the expression profiles of a large number of neuropeptides and neurotransmitters in the same preparation enabled us to conduct the most extensive comparison of the hypothalamic nuclei between zebrafish and mice to date and to reveal several similarities across species.

#### 4.3.1 Zebrafish magnocellular preoptic nucleus (PM), ventral part of the anterior part of the parvocellular preoptic nucleus (PPav), and mouse paraventricular hypothalamic nucleus (Pa)

It has been considered that the teleost PM and mammalian Pa are homologous because they share many features, such as the existence of avp- and oxt-expressing magnocellular neurons (Goodson and Bass, 2001; Olivereau and Olivereau, 1988), innervation by *agrp* and *pomc* neurons (Forlano and Cone, 2007), and the expression of other marker genes such as *nurr1* (Kapsimali *et al*., 2001), *ccka*, *nts*, *penka*, *penkb*, *vipa*, *sst1.1*, *crhb*, and *otpa* (Herget *et al*., 2014). Consistently, the zebrafish PM was clustered closest to the mouse Pa in our analysis. Fifteen of the 17 neuropeptide genes examined showed identical expression status between PM and Pa, suggesting highly similar functionality. Zebrafish PPav also clustered close to the PM. In the brain atlases of some fish species, a portion of the PM is considered to be parvocellular, referred to as the parvocellular part of the PM (PMp), and it is located anteriorly adjacent to the magnocellular part of the PM (PMm) (medaka, Anken and Bourrat, 1998; goldfish, Braford and Northcutt, 1983). In the zebrafish reference atlas (Wullimann *et al*., 1996) it was considered that the PMp was not evident; however, the PPav may correspond to the PMp in other fish species and is closely related to the PM, and therefore, highly similar to Pa in mice.

#### 4.3.2 Zebrafish posterior part of the parvocellular preoptic nucleus (PPp) and mouse medial preoptic area (MPA)

Zebrafish PPp clustered close to the mouse MPA. The preoptic area as a whole is thought to be conserved between teleosts and mammals (Goodson, 2005; O’Connell and Hofmann, 2011); however, the homology of specific nuclei or regions within the preoptic area has not been discussed. The fact that the neuropeptide expression profile of zebrafish PPp was particularly similar to that of mouse MPA may indicate functional similarities between these nuclei. In mammals, MPA harbors estrogen receptor (*Esr1*)-expressing neurons, which play important roles in sexual and parental behaviors (Tsuneoka and Funato, 2021). Although there is no available data on the presence or absence of *esr1* in zebrafish PPp, *esr1* expression in the PPp has been reported in several fish species, such as Atlantic croaker, sea bass, and medaka (Hawkins *et al*., 2005; Muriach *et al*., 2008; Zempo *et al*., 2013). In contrast, *esr1* is also expressed in the PPa and PM in many teleost species. Furthermore, galanin, which is excessively expressed in mammalian MPA (Wu *et al*., 2014; Moffitt *et al*., 2018), was not observed in zebrafish PPp but was abundantly expressed in PPa. Collectively, it may be more accurate to consider that many of the cell populations corresponding to mammalian MPA are located in the PPp in zebrafish, but some are also distributed in other regions of the preoptic area.

#### 4.3.3 Zebrafish dorsal subnucleus of the ventral zone of periventricular hypothalamus (Hvd), anterior tuberal nucleus (ATN), and mouse ventromedial hypothalamic nucleus (VMN)

Zebrafish Hvd and ATN clustered close to the mouse VMN. The similarity between the ATN and VMN has been proposed previously, based on the location, expression of *esr1*, and connections with the preoptic area (Forlano *et al*., 2005; Goodson, 2005; Forlano and Bass, 2011; O’Connell and Hofmann, 2011). Hvd is located medially adjacent to the ATN, but the homologous relationship between Hvd and VMN has not been reported so far. Our data showed that *pdyn*, which is present predominantly in the dorsomedial part of the mouse VMN (Allen Brain Atlas), is strongly expressed in zebrafish Hvd but not in ATN, indicating that Hvd and ATN might represent homologs of different subnuclei in the VMN. Other genes that are highly expressed in the mouse VMN include *Adcyap* and *Bdnf* (Khodai *et al*., 2021). *adcyap1b* is expressed in both the Hvd and ATN in teleosts (Nakamachi *et al*., 2018), and *bndf* is predominantly expressed in the Hvd(Cacialli *et al*., 2016).

Mammalian VMN controls a variety of behavioral and physiological traits such as sexual behavior, aggression, fear response, and energy homeostasis (Silva *et al*., 2013; Hashikawa *et al*., 2017; Khodai *et al*., 2021). In teleost fishes, it has been reported that the Hv and ATN in males are activated by the courtship of competing males (midshipman fish, Petersen *et al*., 2013), and during the change from subordinate to dominant (cichlid fish, Maruska *et al*., 2013), suggesting a role for Hv and ATN in social information processing. Further studies are required to elucidate the degree to which mammalian VMN and fish Hvd/ATN functions are conserved.

## 5. Conclusion

We constructed an expression atlas and database of 38 neuropeptide genes and GABAergic/glutamatergic neuron marker genes in the forebrain of adult zebrafish. By utilizing these data, we proposed subnuclei in the hypothalamic nuclei and compared them with mouse hypothalamic nuclei. This atlas and database will become a useful resource that will contribute to the future development of neuroscience research using zebrafish.

**Supplemental Table 1**. The database of *gal* and *galp* genes in various species was used for phylogenetic tree analysis.

**Supplemental Figure 1.** Phylogenetic tree and synteny analyses of *gal* and *galp* gene families in vertebrates. A: Phylogenetic tree of GAL and GALP amino acid sequences. The accession numbers of the sequences used in this analysis are listed in Supplementary Table 1. Sequences are aligned using ClustalW, and a phylogenetic tree is constructed by the maximum likelihood method using the Molecular Evolutionary Genetics Analysis (MEGA11) software. Numbers at the branch nodes represent the bootstrap confidence level (500 replicates). The scale bar indicates the number of substitutions per site. B: Schematic showing the genes in the genomic area around *gal* and *galp* genes. The arrowhead represents the orientation of the gene. Orthologous genes among vertebrate species are indicated in the same color.

**Supplemental Figure 2.** Summary of individual differences in neuropeptide expression. Photographs of four individual fish are shown for each gene for which individual differences in expression are observed. Some sections are subjected to methyl green nuclear staining. Scale bar = 100 μm.

## Abbreviations

Zebrafish brain regions and nuclei: 
A: anterior thalamic nucleus
APN: accessory pretectal nucleus
ATN: anterior tuberal nucleus
CIL: central nucleus of the inferior lobe
CM: corpus mamillare
CP: central posterior thalamic nucleus
CPN: central pretectal nucleus
Cans: commissura ansulata
Cantv: commissura anterior, pars ventralis
Ce: cerebellum
Cpop: commissura postoptica
D: dorsal telencephalic area
DAO: dorsal accessory optic nucleus
DIL: diffuse nucleus of the inferior lobe
DOT: dorsomedial optic tract
DP: dorsal posterior thalamic nucleus
DTN: dorsal tegmental nucleus
Dc: central zone of D
Dd: dorsal zone of D
DiV: diencephalic ventricle
Dl: lateral zone of D
Dld: dorsal part of Dl
Dlv: ventral part of Dl
Dm: medial zone of D
Dma: anterior part of Dm
Dmp: posterior part of Dm
Dp: posterior zone of D
E: epiphysis
ECL: external cellular layer of olfactory bulb including mitral cells
ENd: dorsal part of entopeduncular nucleus
ENv: ventral part of entopeduncular nucleus
EW: Edinger-Westphal nucleus
GL: glomerular layer of olfactory bulb
Had: dorsal habenular nucleus
Hadl: lateral part of Had
Hadm: medial part of Had
Hav: ventral habenular nucleus
Hc: caudal zone of periventricular hypothalamus
Hcad: anterodorsal part of Hc
Hcai: anterointermediate part of Hc
Hcav: anteroventral part of Hc
Hcpd: posterodorsal part of Hc
Hcpi: posterointermediate part of Hc
Hcpv: posteroventral part of Hc
Hd: dorsal zone of periventricular hypothalamus
Hddl: dorsolateral part of Hd
Hdm: medial Hd
Hdvl: ventrolateral part of Hd
Hv: ventral zone of periventricular hypothalamus
Hvd: dorsal part of Hv
Hvv: ventral Hv
Hyp: hypothalamus
I: intermediate thalamic nucleus
IC: intercalated nucleus
ICL: internal cellular layer of olfactory bulb
III: oculomotor nerve
IN: intermediate nucleus
LH: lateral hypothalamic nucleus
LR: lateral recess
MNV: mesencephalic nucleus of trigeminal nerve
Med: medulla oblongata
NIII: oculomotor nucleus
NLL: nucleus of the lateral lemniscus
NLT: nucleus lateralis tuberis
NMLF: nucleus of medial longitudinal fascicle
NT: nucleus taeniae
OB: olfactory bulb
OpT: optic tectum
P: posterior thalamic nucleus
PCN: paracommissural nucleus
PGZ: periventricular gray zone of optic tectum
PGa: anterior preglomerular nucleus
PGc: caudal preglomerular nucleus
PGl: lateral preglomerular nucleus
PGm: medial preglomerular nucleus
PL: perilemniscal nucleus
PM: magnocellular preoptic nucleus
PO: posterior pretectal nucleus
POF: primary olfactory fiber layer
PPa: anterior part of parvocellular preoptic nucleus
PPad: dorsal part of PPa
PPav: ventral part of PPa
PPd: dorsal part of periventricular pretectal nucleus
PPp: posterior part of parvocellular preoptic nucleus
PPv: ventral part of periventricular pretectal nucleus
PR: posterior recess
PSm: magnocellular superficial pretectal nucleus
PSp: parvocellular superficial pretectal nucleus
PTN: posterior tuberal nucleus
PVO: paraventricular organ
R: rostrolateral nucleus
RT: rostral tegmental nucleus
SC: suprachiasmatic nucleus
SG: subglomerular nucleus
SD: saccus dorsalis
TGN: tertiary gustatory nucleus
TL: torus longitudinalis
TLa: torus lateralis
TPp: periventricular nucleus of posterior tuberculum
TS: torus semicircularis
TSc: central nucleus of torus semicircularis
TSvl: ventrolateral nucleus of torus semicircularis
TeO: optic tectum
Tel: telencephalon
V: ventral telencephalic area
VAO: ventral accessory optic nucleus
VL: ventrolateral thalamic nucleus
VM: ventromedial thalamic nucleus
VOT: ventrolateral optic tract
Val: lateral division of valvula cerebelli
Vam: medial division of valvula cerebelli
Vas: vascular lacuna of area postrema
Vc: central nucleus of V
Vd: dorsal nucleus of V
Vi: intermediate nucleus of V
Vl: lateral nucleus of V
Vp: postcommissural nucleus of V
Vs: supracommissural nucleus of V
Vv: ventral nucleus of V
ZL: zona limitans
Mouse brain regions and nuclei: 
AH: anterior hypothalamic nucleus
Arc: arcuate hypothalamic nucleus
AVPe: anteroventral periventricular nucleus
DM: dorsomedial nucleus of the hypothalamus
d/vPM: dorsal and ventral premammillary nucleus
LHA: lateral hypothalamic area
LPO: lateral preoptic area
MB: medial preoptic nucleus
MnPO: median preoptic nucleus
MPA: medial preoptic area
MPO: medial preoptic nucleus
Mtu: medial tuberal nucleus
Pa: paraventricular hypothalamic nucleus
Pe: periventricular hypothalamic nucleus
PH: posterior hypothalamic nucleus
PS: parastrial nucleus
PSTh: parasubthalamic nucleus
PVp: periventricular hypothalamic nucleus, posterior part
Rch: retrochiasmatic area
SCh: suprachiasmatic nucleus
SCO: subcommisural organ
SO: supraoptic nucleus
Spa: subparaventricular zone
VLPO: ventrolateral preoptic nucleus
VMN: ventromedial hypothalamic nucleus

## Acknowledgments

We thank the RIKEN Research Resources Division for providing services and equipment. We thank Nobuko Mataga for her support and coordination of the collaborative research with the RIKEN Support Unit for Bio-Material Analysis. We thank Miwa Masuda for providing the protocol for *in situ* hybridization; members of Yoshihara Lab for critical discussions; Shin-ichi Higashijima for providing plasmids for *slc17a6a*, *slc17a6b*, and *slc17a7a*; and Chigaya Ito and Satomi Yamashita for taking good care of zebrafish. The present work was supported by JSPS KAKENHI Grant Number 15H06860, 19K16304 (to T. H-K.), RIKEN Special postdoctoral researcher budget (to T. H-K.), Funding support for researchers returning to work/research from interruption from RIKEN Diversity Promotion Office (to T H-K). This work was also supported in part by Grants-in-Aid for Scientific Research in the Innovative Area “LipoQuality” (18H04683 to Y.Y.) from the Ministry of Education, Culture, Sports, Science and Technology of Japan, and grants from Ono Medical Research Foundation (to Y.Y.), Kao Corporation (to Y.Y.), and JST NBDC Grant Number JPMJND2201 (to S.O.).

