## Supplemental Table 1, Supplemental Figures 1, 2 for "An atlas and database of neuropeptide gene expression in the adult zebrafish forebrain"

Supplemental Table 1
Accession numbers of sequences used in phylogenetic tree analysis.

| Species | Peptide name | Chromosome/scaffold | GenBank accession no. | Amino acid size |
|---|---|---|---|---|
| Spotted gar (*Lepisosteus oculatus*) | GAL | LG27 | XP_015193576.1 | 142 |
| Reedfish (*Erpetoichthys calabaricus*) | GAL | Chr. 2 | XP_028651983.1 | 142 |
| Reedfish (*Erpetoichthys calabaricus*) | GALP | NW_021316950.1 | XP_028647327.1 | 119 |
| Zebrafish (*Danio rerio*) | GAL | Chr. 25 | NP_001333168.1 | 142 |
| Zebrafish (*Danio rerio*) | GALP | Chr. 12 | NP_001314905.1 | 114 |
| Channel catfish (*Ictalurus punctatus*) | GAL | Chr. 8 | XP_017329224.1 | 142 |
| Channel catfish (*Ictalurus punctatus*) | GALP | Chr. 13 | XP_017339903.1 | 119 |
| Brown trout (*Salmo trutta*) | GAL | Chr. 7 | XP_029613854.1 | 142 |
| Brown trout (*Salmo trutta*) | GALP | Chr. 10 | XP_029620010.1 | 117 |
| Nile tilapia (*Oreochromis niloticus*) | GAL | LG7 | XP_025764384.1 | 142 |
| Japanese medaka (*Oryzias latipes*) | GAL | Chr. 6 | XP_011471141.1 | 142 |
| Torafugu (*Takifugu rubripes*) | GAL | NW_021821634.1 | XP_029687971.1 | 145 |
| Tropical clawed frog (*Xenopus tropicalis*) | GAL | Chr. 4 | XP_017948725.2 | 146 |
| Tropical clawed frog (*Xenopus tropicalis*) | GALP | Chr. 7 | XP_004916350.1 | 119 |
| Green anole (*Anolis carolinensis*) | GAL | Chr. 1 | XP_008106931.1 | 140 |
| Green anole (*Anolis carolinensis*) | GALP | NW_003338902.1 | XP_016852185.1 | 120 |
| Australian saltwater crocodile (*Crocodylus porosus*) | GAL | NW_017728937.1 | XP_019402803.1 | 141 |
| Australian saltwater crocodile (*Crocodylus porosus*) | GALP | NW_017728947.1 | XP_019406661.1 | 116 |
| Chicken (*Gallus gallus*) | GAL | Chr. 5 | NP_001153150.1 | 141 |
| Mouse (*Mus Musculus*) | GAL | Chr. 19 | NP_001316596.1 | 141 |
| Mouse (*Mus Musculus*) | GALP | Chr. 7 | NP_821171.1 | 117 |
| Human (*Homo sapiens*) | GAL | Chr. 11 | NP_057057.2 | 123 |
| Human (*Homo sapiens*) | GALP | Chr. 19 | NP_149097.1 | 116 |

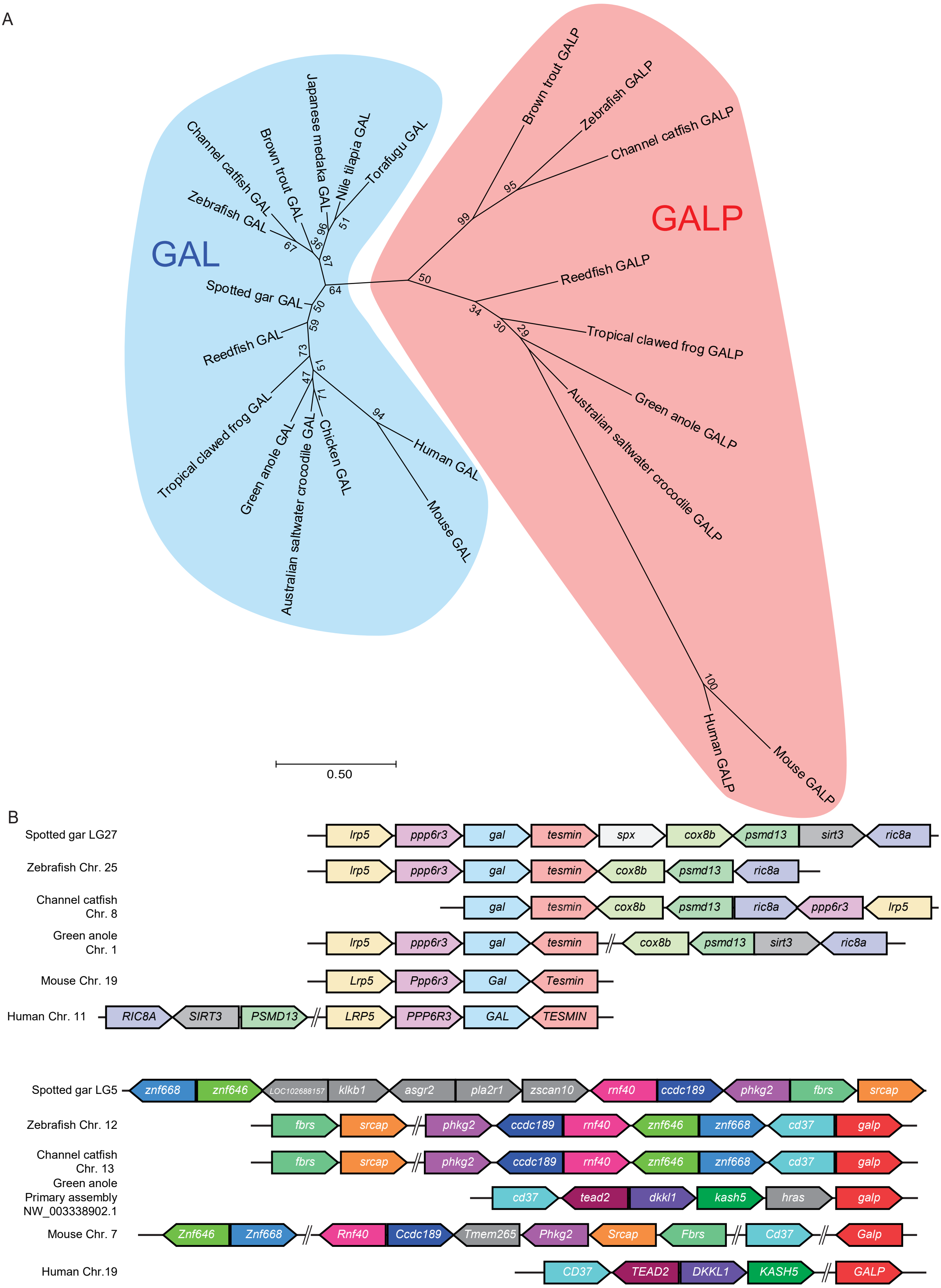
A
GAL
GALP
Channel catfish GAL
Brown trout GAL
Japanese medaka GAL
Nile tilapia GAL
Torafugu GAL
Zebrafish GAL
Spotted gar GAL
Reedfish GAL
Tropical clawed frog GAL
Green anole GAL
Australian saltwater crocodile GAL
Chicken GAL
Human GAL
Mouse GAL
Brown trout GALP
Zebrafish GALP
Channel catfish GALP
Reedfish GALP
Tropical clawed frog GALP
Green anole GALP
Australian saltwater crocodile GALP
Human GALP
Mouse GALP
0.50
B
Spotted gar LG27: lrp5, ppp6r3, gal, tesmin, spx, cox8b, psmd13, sirt3, ric8a
Zebrafish Chr. 25: lrp5, ppp6r3, gal, tesmin, cox8b, psmd13, ric8a
Channel catfish Chr. 8: gal, tesmin, cox8b, psmd13, ric8a, ppp6r3, lrp5
Green anole Chr. 1: lrp5, ppp6r3, gal, tesmin, cox8b, psmd13, sirt3, ric8a
Mouse Chr. 19: Lrp5, Ppp6r3, Gal, Tesmin
Human Chr. 11: RIC8A, SIRT3, PSMD13, LRP5, PPP6R3, GAL, TESMIN
Spotted gar LG5: znf668, znf646, LOC102688157, klkb1, asgr2, pla2r1, zscan10, rnf40, ccdc189, phkg2, fbrs, srcap
Zebrafish Chr. 12: fbrs, srcap, phkg2, ccdc189, rnf40, znf646, znf668, cd37, galp
Channel catfish Chr. 13: fbrs, srcap, phkg2, ccdc189, rnf40, znf646, znf668, cd37, galp
Green anole Primary assembly NW_003338902.1: cd37, tead2, dkkl1, kash5, hras, galp
Mouse Chr. 7: Znf646, Znf668, Rnf40, Ccdc189, Tmem265, Phkg2, Srcap, Fbrs, Cd37, Galp
Human Chr.19: CD37, TEAD2, DKKL1, KASH5, GALP

Supplemental Figure 1

Phylogenetic tree and synteny analyses of *gal* and *galp* genes in vertebrates. A: A phylogenetic tree of GAL and GALP amino acid sequences. The GenBank accession numbers of sequences used in the analysis are listed in Supplemental Table 1. Sequences were aligned by clustalW and the phylogenetic tree was constructed by the maximum likelihood method using MEGA11 software. Numbers at branch nodes represent the bootstrap confidence level (500 replications). The scale bar shows the number of substitutions per site. B: A schematic showing syntenic analysis of genes around *gal* and *galp* loci. The arrowheads represent the orientation of the genes. Orthologous genes among vertebrate species are indicated in the same color.

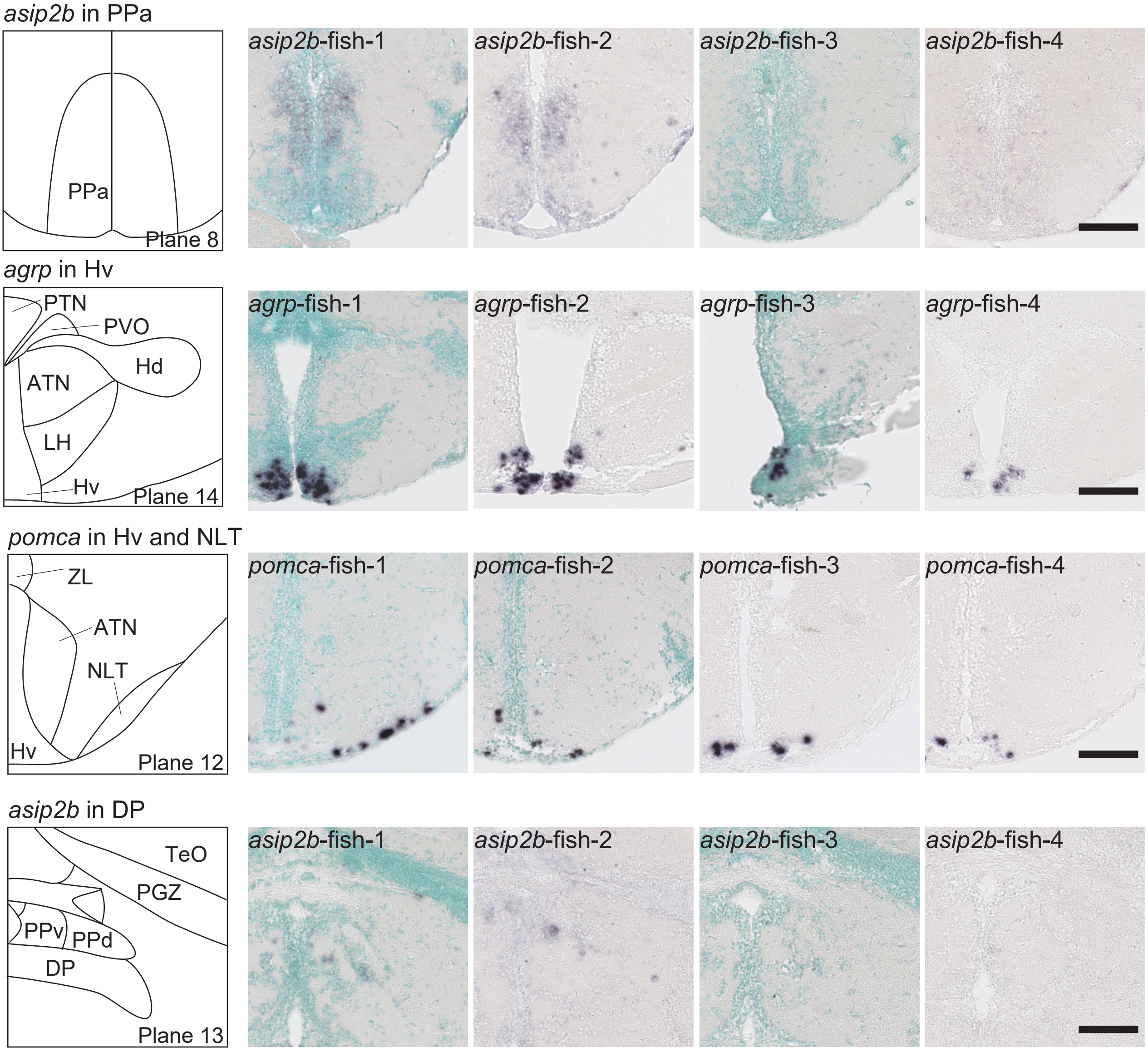


Supplemental Figure 2

Summary of individual differences in the neuropeptide expression. For each of the genes that displayed individual differences in expression, microscopic images of four individual fish are shown. Several images include nuclear counterstaining with methyl green and the others do not. Scale bar = 100 μm.
